# The TCRs of pathogenic Th17 cells in arthritogenic mice are shifted toward a Treg-like repertoire

**DOI:** 10.1101/2024.03.01.582868

**Authors:** Mara A. Llamas-Covarrubias, Atsushi Tanaka, Martin Loza, Diego Diez, Zichang Xu, Ee Lyn Lim, Shunsuke Teraguchi, Shimon Sakaguchi, Daron M. Standley

**Author notes:** **Correspondence:** Daron M. Standley. 3-1 Yamadaoka, Suita, Osaka 565-0871, Japan. **Correspondence:** Mara A. Llamas-Covarrubias. 3-1 Yamadaoka, Suita, Osaka 565-0871, Japan.

## Abstract

Mutations in the ZAP-70 gene that cause moderate attenuation of T cell receptor (TCR) signaling in mice can result in autoimmune manifestations. One explanation for this pathology is a shift in the regulatory-conventional (Treg-Tconv) T cell repertoire composition. To test this hypothesis, we characterized the single-cell gene expression profiles and TCR repertoires of Tconv and Treg CD4+ T cells of arthritic (ZAC), poised (SKG) ZAP-70 mutant, and wild-type (WT) mice. We identified a group of Th17 cells which exhibited a pathogenic signature and occurred exclusively in inflamed joints of ZAC mice. Such pathogenic signature was uniquely detected in CD4+ T cells obtained from inflamed joints of RA patients. Overall, the Tconv repertoires of ZAP-70 mutant mice were increasingly similar to the repertoires of WT Tregs, and this effect was most notable in the subset of pathogenic Th17 cells. Our results support a model where, upon moderate ZAP-70-mediated signal weakening, T cells that would normally develop into Tregs, instead develop into self-reactive Tconvs, resulting in a breakdown in self-tolerance and susceptibility to autoimmune arthritis.

## Introduction

Autoimmune diseases encompass a diverse group of disorders characterized by aberrant T and B cell responses against the body’s own tissues. These conditions are a major source of morbidity and disability worldwide, disproportionately affecting young women(1). Despite extensive research, the molecular mechanisms driving the development and activation of self-reactive immune cells remain poorly understood. Previous studies have implicated a range of factors in autoimmune disease pathogenesis, including viral infections(2), X chromosome-linked ribonucleoproteins (3), and the HLA presentation of misfolded proteins (4), among others. Independent of these models, our group recently demonstrated that a moderate, systemic reduction in the T cell receptor (TCR) signaling protein ZAP-70 results in autoimmune arthritis and colitis in mice (5), while more severe impairment has been shown to lead to immunodeficiency (6).

The link between weakened TCR signaling and autoimmune disease appears to hinge on TCR repertoire composition rather than signaling intensity in the periphery. During T cell maturation, the repertoire is shaped by thymic selection, a process that ensures T cells are both functional (MHC-restricted, through positive selection) and self-tolerant (through negative selection). High-affinity interactions between TCRs and self-peptide-MHC complexes typically lead to either clonal deletion or selection into the Treg population (7). However, a systemic weakening of TCR signaling could disrupt this thymic selection, allowing T cells with moderately to strongly self-reactive TCRs to evade negative selection and enter the mature conventional T cell (Tconv) repertoire. This concept, formally known as the “Repertoire Shift” hypothesis, is illustrated in Fig. 1a.

**Figure 1.**
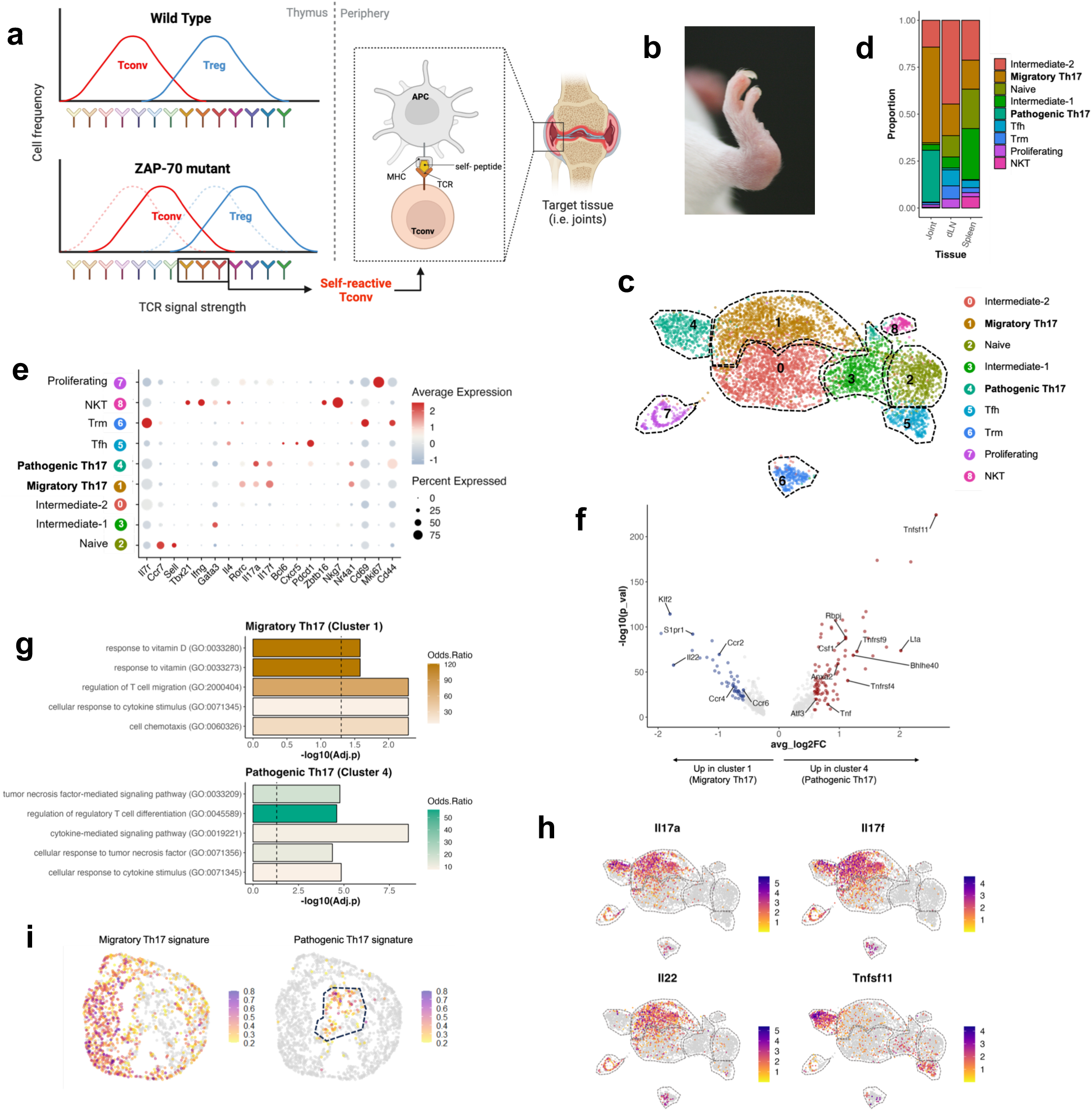
Two transcriptionally different Th17 cell populations exist in inflamed joints of arthritic ZAC mice. a) **a)** Normally, during T cell development in thymus, high-intensity interactions between a TCR and self-pMHC will induce either clonal deletion or selection into the Treg repertoire (upper plot); however, a systemic weakening of TCR signaling could disrupt this mechanism by allowing T cells with moderately to strongly self-reactive TCRs to evade negative selection and enter the mature conventional Tconv repertoire (lower plot). These cells would later be activated by self-peptides in the periphery, accumulate in target tissues and orchestrate tissue damage. **b)** Lateral view of a hindpaw of a 23-week old arthritic ZAC mouse. **c)** UMAP plot of Tconv cells obtained from joints, draining lymph node (dLN) and spleen of arthritic ZAC mice colored by cluster. **d)** Proportion of annotated Tconv cell populations in tissues of arthritic ZAC mice. **e)** Relative expression of canonical Tconv markers across gene expression clusters. Circle color and intensity represent the average expression whereas circle size represents the percentage of cells from each cluster expressing the gene. **f)** Volcano plot of the differentially expressed genes (DEG) among cluster 1 and cluster 4. **g)** Top 5 gene Ontology (GO) Terms enriched in DEGs from cluster 1 vs cluster 4. The color intensity of bars represents the odds ratio (OR) of enrichment. **h)** UMAP plot of Tconv cells from tissues of arthritic ZAC mice colored according to their expression of *Il17a*, *Il17f*, *Il22* or *Tnfsf11*. **i)** Module scores for the gene signatures of migratory Th17 and pathogenic Th17 cells in a public scRNAseq dataset of CD4^+^ T cells from inflamed synovia of RA patients (n=3)(18). Created with BioRender.com

While the “Repertoire Shift” hypothesis aligns with previous observations in ZAP-70 mutant mice (5, 8, 9), it has not been rigorously tested. Advances in repertoire sequencing and single-cell resolution technologies have now enabled the first direct test of this hypothesis. To investigate it, we analyzed the transcriptome and immune repertoires of CD4+ T cells from wild-type (WT), and two ZAP-70 mutant mouse models: non-arthritic SKG, and arthritic ZAC, at the single-cell level. Our findings revealed distinct migratory and pathogenic populations of Th17 cells in arthritic mice. Notably, these pathogenic cells exhibited high expression of osteoclastogenic genes, were localized specifically to inflamed joints, and were absent in both WT and SKG mice. Strikingly, a population with a similar pathogenic signature was uniquely identified in T cells from human RA joints. Moreover, the TCR repertoire of these pathogenic Th17 cells was unique and convergent. Overall, we found an increased similarity of ZAP-70 Tconv repertoires to the repertoires of large public WT Treg datasets, as demonstrated by two independent metrics. In addition, the Treg-like nature of ZAP-70 mutant Tconv TCRs was most notable in the subset of pathogenic Th17 cells. Additionally, the overall similarity between Treg and Tconv repertoires was significantly diminished in arthritic mice, suggesting that autoimmunity may result from a depletion of critical Treg populations and the expansion of self-reactive Tconvs. Our findings suggest that restoring the balance between Treg and Tconv repertoires could be a promising therapeutic strategy for treating autoimmune diseases.

## Results

### Arthritic ZAC mice harbor two groups of Th17 cells with distinct gene programs in their joints

The ZAP-70 mutant mice (ZAC, SKG) used in this study are characterized by the spontaneous development of autoimmune arthritis (Fig. 1b), with disease progression typically accelerated in the SKG model through the stimulation of innate immunity(5, 8, 9). Unlike other widely used mouse models of arthritis, such as collagen-induced arthritis (CIA)(10), the arthritis in ZAP-70 mutant mice is genuinely autoimmune, mediated primarily by T-cells, and closely resembles human RA(5, 8, 9). Therefore, ZAP-70 mutant mouse models are particularly well-suited for investigating self-reactive TCR repertoires. To study the transcriptomes and immune repertoires of self-reactive Tconv (CD3^+^CD4^+^Foxp3^-^) cells in-depth, we sorted cells from the joints, draining lymph nodes (dLN), and spleens of arthritic ZAC mice and performed RNA+V(D)J single-cell sequencing. Gene expression analysis revealed a considerable overlap in the transcriptional profiles of spleens and dLNs but a distinct profile for joint cells (Supplementary Fig. 1a). Unsupervised clustering delineated nine gene expression clusters, including two (clusters 1 and 4) expressing canonical Th17 markers (*Rorc, Il17a, Il17f*) but with different gene expression programs and tissue abundances (Supplementary Note 1, Fig. 1c-e, Supplementary Fig. 1b-d). Cluster 1 was abundant in joints, but was also detected in the spleen and dLNs. Relative to cluster 4, cells in cluster 1 differentially expressed *Klf2*, *S1pr1* and chemokine receptors, and it was enriched in the regulation of T cell migration (GO: 20000404) and cell chemotaxis (GO: 0060326) gene ontology (GO) Terms (Fig. 1f-g, Supplementary Table 4-5) suggesting a phenotype of migratory cells. In contrast, cluster 4 was almost exclusively observed in joints, and highly expressed *Tnfsf11* (encoding for RANKL) along with other osteoclastogenic genes (*Csf1*, *Anxa2*), and overall, the osteoclast differentiation (GO:0030316) GO term was significantly enriched (Fig. 1d, f-g, Supplementary Table 5). Moreover, this cluster uniquely expressed transcription factors associated with a pathogenic Th17 phenotype: *Rbpj*(11)*, Crem*(12)*, Atf3*(13)*, Bhlhe40*(14), and *Igfbp7*(15). These results indicate that cluster 4 contains pathogenic Th17 cells with an increased osteoclastogenic potential. Thus, we annotated clusters 1 and 4 as migratory and pathogenic Th17, respectively. Interestingly, cluster 1 and cluster 4 could be clearly distinguished by their differential expression of *Il22* or *Tnfsf11* (Fig. 1h). Trajectory inference of the different Tconv cell states resulted in the identification of the pathogenic Th17 cluster as one of the endpoint lineage states (Supplementary Fig. 1e), consistent with their effector phenotype. Of note, we did not detect clear and specific expression of marker genes of previously reported populations of pathogenic Th17 cells (Th17.1(16), exFoxP3(17)) in cluster 4, suggesting that it represents a distinct group of pathogenic cells (Supplementary Fig. 1f-g).

To contextualize our findings in arthritic ZAC mice, we replicated the analyses in WT and non-arthritic SKG mice. Here, we sampled Tconv from their spleens and integrated the ZAC spleen data in order to facilitate direct comparison of clusters (Supplementary Note 2, Supplementary Fig. 2a-g). This approach revealed that the general Th17 cell abundance increased progressively from WT to SKG to ZAC mice (Supplementary Fig. 2d). To analyze whether the transcriptional programs of these Th17 cells were similar to those in the migratory and pathogenic Th17 cells, we calculated gene module scores for their signature genes (Supplementary Table 6). Expression of the migratory Th17 signature was clearly detected and gradually increased from WT, to SKG to ZAC splenic Th17 cells; however, the pathogenic Th17 signature was largely undetected (Supplementary Fig. 3a-b).

Thus, we demonstrated the presence of two distinct groups of Th17 cells in arthritic ZAC mice: migratory Th17 cells, expressing *Il22*, and pathogenic Th17 cells, which highly express *Tnfsf11* and uniquely localize within inflamed joints.

### The transcriptional signature of pathogenic Th17 cells is present in inflamed joints of human RA but not in other autoimmune diseases or non-autoimmune arthritis

To investigate the presence of cells resembling pathogenic Th17 cells in human tissues, we calculated migratory and pathogenic Th17 gene signature module scores in publicly available single-cell RNAseq datasets. We analyzed data obtained from three sources: inflamed synovia(18) and PBMCs(19) of RA patients; synovia and infrapatellar fat pad (IPFP) of osteoarthritis (OA) patients(20); peripheral CD4+ T cells from healthy controls and patients with other autoimmune diseases(21). Detailed descriptions of the datasets are provided in Supplementary Table 7.

Our analysis revealed that the migratory Th17 signature was clearly detected in all datasets (Fig. 1i, Supplementary Fig. 3c-e). In contrast, expression of the pathogenic Th17 signature was nearly undetectable in all samples (Supplementary Fig. 3c-e) except in CD4^+^ T cells derived from inflamed joints of RA patients (Fig. 1i); in addition, its expression was non-overlapping with the expression of the migratory Th17 signature, mirroring our observations in ZAC joints. Notably, as in ZAC joints, the expression of Th17.1 or exFoxP3 markers was not specifically detected in the cluster of cells expressing the pathogenic Th17 gene signature in human RA joints (Supplementary Fig. 4a-d). Since OA is a form of non-autoimmune arthritis(22), our results indicate that cells with a pathogenic Th17 transcriptional signature are likely to be specifically involved in the pathogenesis of human autoimmune arthritis.

### Pathogenic Th17 cells are expanded, convergent, and contain public clones

We next asked whether the TCR repertoires of the two Th17 clusters were also distinct. First, we examined clonal expansion and found that large and medium clones were abundant in both Th17 cell groups (Fig. 2a, Supplementary Fig. 5a-b). Next, we evaluated the repertoire similarity of both Th17 groups to other Tconv populations and between each other by using the Morisita-Horn index. At the exact paired CDR3 amino acid sequence level, the repertoire of migratory Th17 cells was largely similar to that of other Tconv repertoires from ZAC tissues (Fig. 2b). In contrast, the pathogenic Th17 repertoire was less similar to other Tconv cells, and the similarity between the two Th17 cell repertoires was rather low (Fig. 2b). These results suggest that pathogenic Th17 cells possess a unique repertoire, presumably due to the targeting of a specific set of antigens.

**Figure 2.**
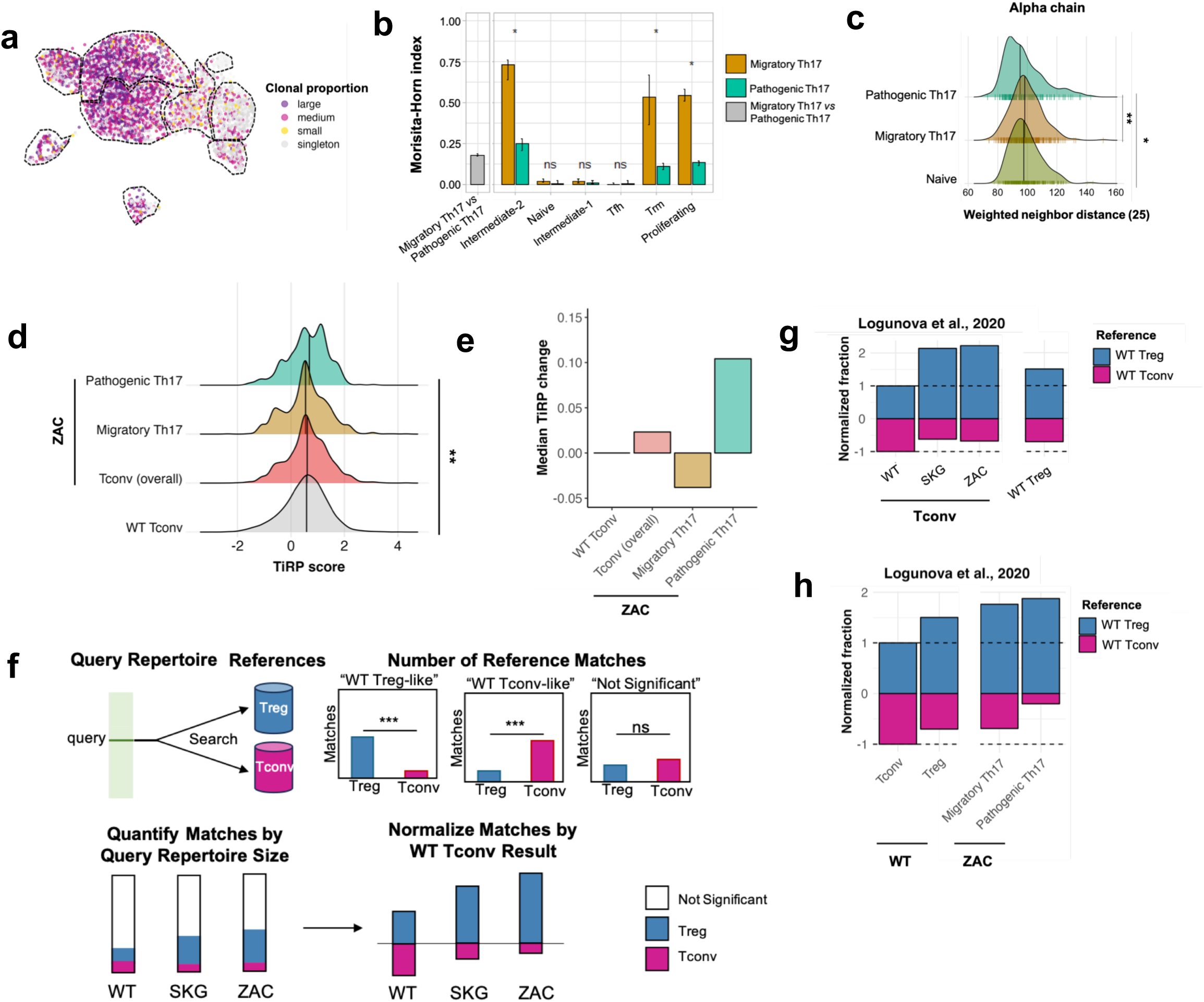
The TCR repertoire of pathogenic Th17 cells is unique, convergent, and self-reactive. **a)** UMAP plot of Tconv cells from tissues of arthritic ZAC mice, colored according to clonal proportion classification. **b)** Morisita-Horn index of the paired CDR3 amino acid similarity between the migratory and pathogenic Th17 repertoires (gray bar); and between both Th17 populations vs the other Tconv phenotypes in ZAC tissues (brown and green bars). **c)** Distribution of neighbor distances to the nearest 25th percentile (methods) in the alpha chain repertoires of naïve, migratory Th17 and pathogenic Th17 cells. **d)** Distribution of TiRP scores for WT Tconv, and overall Tconv, migratory Th17, and pathogenicTh17 repertoires of ZAC mice. **e)** Median TiRP score change of overall Tconv, migratory Th17, and pathogenicTh17 repertoires of ZAC mice relative to WT Tconv. **f)** Description of the method employed for the assessment of repertoire similarity to reference WT Treg and WT Tconv repertoires. **(g-h)** Normalized fraction of query WT, SKG and ZAC Tconv repertoires **(g)** or WT Tconv, WT Treg, ZAC migratory and pathogenic Th17 repertoires **(h)** matching with reference WT Tconv (magenta section) or WT Treg repertoires (blue section) obtained from a representative reference dataset(25). *: p<0.05; **:p<0.01; Wilcoxon Test.

We next evaluated intra-repertoire TCR similarity as a measure of repertoire convergence. For this purpose, we used TCRdist(23), which computes the distance between the CDR1, CDR2 and CDR3 amino acid sequences of single or paired TCR chains. Using TCRdist, we independently calculated distances among all receptors in the two Th17 clusters, as well as the naïve cluster for reference. Because a bias in the distribution of the distances to lower values indicates higher similarity within the receptors, it can be interpreted as an indication of repertoire convergence. We found that the TCR distance distribution of the pathogenic Th17 repertoire trended lower in paired and single chain analysis, especially for the alpha chain (Fig. 2c, Supplementary Fig. 5c). This result indicates that the repertoire of pathogenic Th17 cells is more convergent than that of migratory Th17 cells. Finally, we identified a total of 9 unique clones shared in two or more individuals (public clones) and these included expanded clones within the Th17 compartments (Supplementary Fig. 5d, Supplementary Table 8). Together, these results indicate that although the Th17 response in arthritic ZAC mice is polyclonal, the repertoire of tissue-specific pathogenic Th17 cells is unique, convergent, and public. Given their specific location in joints, and exclusive similarity to Th17 T cells in human RA, our results suggest that such pathogenic Th17 cells may recognize a restricted set of self-antigens in the affected tissues.

### TCR repertoires of pathogenic Th17 cells are more Treg-like than Tconv-like

The repertoire shift hypothesis postulates that ZAP-70 hypomorphic mutations would lead to the appearance of TCR repertoires in Tconvs that would typically be deleted or found in WT Tregs(5, 9) (Fig. 1a). To test this prediction, we used a machine learning model, TiRP (TCR-intrinsic regulatory potential), that measures the intrinsic Treg propensity of a given TCR, based on sequence features(24). It has previously been shown that self-reactive TCRs typically have higher TiRP scores than non self-reactive TCRs(24). Consistent with these reports, our experimentally determined Treg datasets exhibited higher TiRP scores than WT Tconv cells, and their respective Tconv compartments (Supplementary Fig. 6a-b). In contrast, the ZAP-70 mutant Tconvs exhibited significantly higher TiRP scores than WT Tconvs or even WT Tregs (Supplementary Fig. 6a-b). Furthermore, we found that pathogenic Th17 TiRP scores were significantly higher than those of WT Tconvs (Fig. 2d-e). These finding support a model where TCRs that would normally be deleted or develop into Tregs instead develop into pathogenic Tconvs upon moderate TCR signal weakening. We next focused on the Treg and Tconv TCR sequences that have been identified in previous studies (Supplementary Table 10) (25–28) in order to confirm whether or not we could observe similarities to the TCRs in our study. To this end, we treated each of our experimentally-determined (WT, SKG and ZAC Tconv and Treg) TCRs as a “query” that was used to search reference WT Tconv or WT Treg datasets (Logunova(25), Lu(26), Ko(27), Wolf(28)). For each query, we computed the number of reference WT Tconv and WT Treg sequence matches to CDR1, CDR2 and CDR3. The number of matches (normalized by the total size of each reference dataset) was deemed “Treg-like”, “Tconv-like” or “not significant” using a hypergeometric p-value cutoff of 0.001 (Fig. 2f). Considering the largest (Logunova(25)) reference dataset, we found that the number of Treg-like TCRs within the SKG and ZAC Tconv T cells was approximately twice as high as that within WT Tconvs, and was higher even than that of query WT Tregs (Fig. 2g). In addition, the fraction of Tconv-like TCRs was notably reduced within ZAP-70 mutant Tconv queries (Fig. 2h). Similar trends were observed for the other reference datasets (Supplementary Fig. 6c). Significantly, the ZAC pathogenic Th17 repertoire exhibited the highest fraction of Treg-like TCRs along with the smallest fraction of Tconv-like TCRs (Fig. 2h, Supplementary Fig. 6d), indicating that the TCRs of pathogenic Th17 cells were largely absent from WT Tconv repertoires. Of note, pathogenic Th17 cells from ZAC mice did not show any significant WT Treg or WT Tconv matches within the Ko et al., 2020(27) dataset (Supplementary Fig. 6d). These observations support TiRP predictions that ZAP-70 mutant Tconv repertoires are more Treg-like than Tconv-like, and that this trend is most pronounced in the pathogenic Th17 cells.

### Evidence of missing TCR specificities in ZAP-70 mutant WT Treg repertoires

A second consequence of weakening of TCR signaling is that it generates “holes” in the normal Treg repertoire. It has been reported that missing specificities in the Treg repertoire have a large impact on the suppressive function against self-reactive T cells(29, 30). This raises the possibility that an absence of Treg specificities within ZAP-70 mutant mice repertoires contribute to a loss in tolerance toward specific self-antigens. To explore this hypothesis, we analyzed the Treg-Tconv repertoire similarity within each mouse and compared it across groups (WT, SKG, ZAC). The Morisita-Horn index for exact CDR3 overlap showed a notably reduced Treg-Tconv similarity in ZAC mice relative to WT or SKG, especially at the single chain level (Fig. 3a). The Treg-Tconv overlap was generally distributed across effector phenotypes (Tconv: intermediate, Th17 and Tfh. Treg: intermediate and effector) in non-arthritic mice (Supplementary Fig. 7a-h), suggesting a suppressive potential for shared Tregs towards activated Tconvs; however, such enrichment was not observed in ZAC mice. To further test these observations, we again employed TCRdist(23) and calculated repertoire-repertoire correlations. We found that WT Treg and WT Tconv repertoires were well correlated, whereas SKG and ZAC Tconv repertoires exhibited lower correlation with their Treg repertoires. Indeed, the ZAC Treg repertoire was poorly correlated with other groups in general, indicating a reduced similarity to any other tested repertoires (Fig. 3b). Together, these results are consistent with a scenario wherein ZAP-70 mutant were unable to select certain Treg TCR specificities normally present in WT mice and capable of suppressing autoimmune arthritis. This, along with a more self-reactive Th17 population, contributes to a loss of tolerance and the development of autoimmune arthritis.

**Figure 3.**
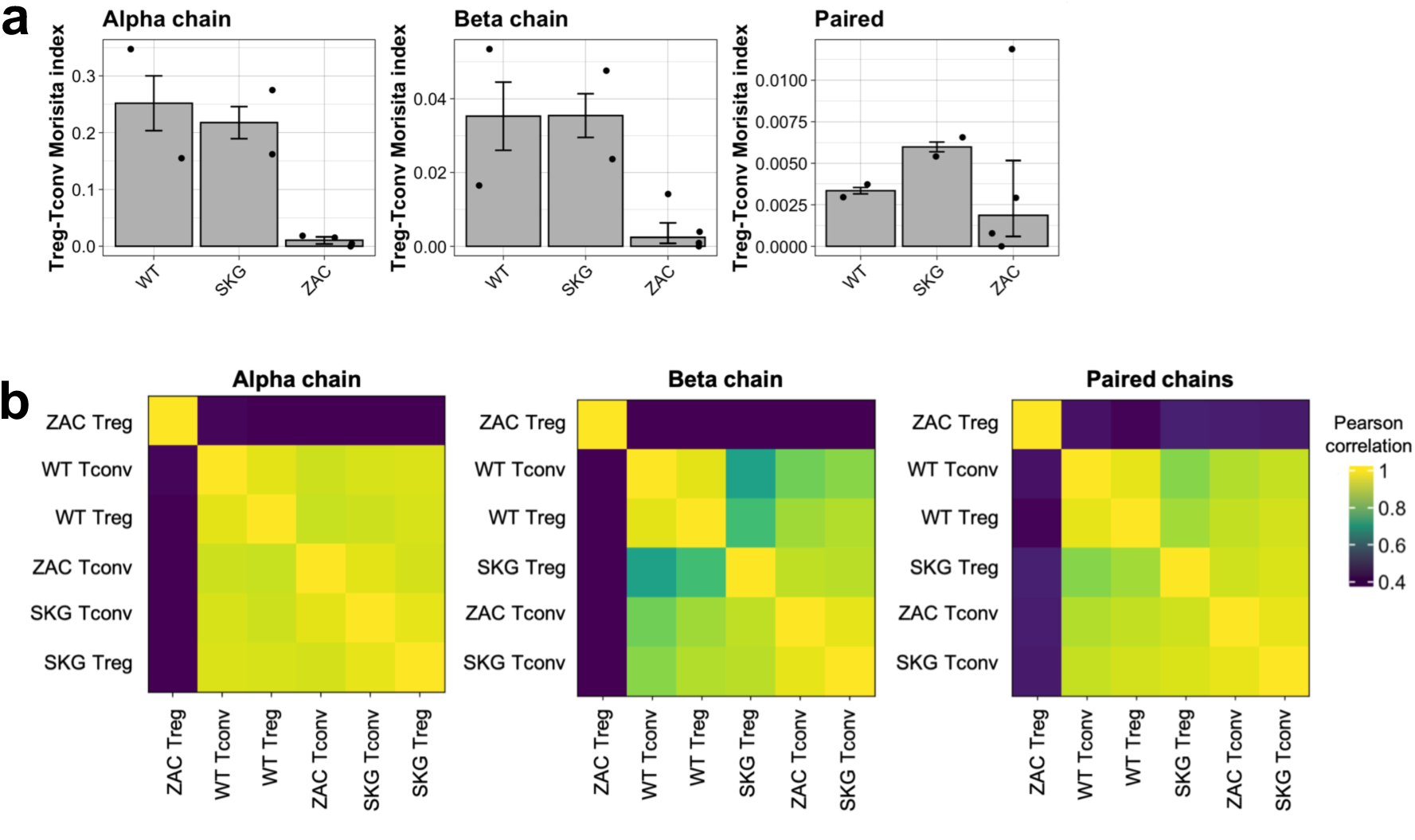
Intra-Treg-Tconv similarity in WT, SKG and ZAC repertoires. **a)** Morisita-Horn index calculated at the exact CDR3 amino acid level between the Tconv and Treg compartments of WT, SKG and ZAC mice. **b)** Heatmap of weighted neighbor distance correlations between WT, SKG and ZAC Treg and Tconv repertoires as calculated by TCRdist (methods), higher correlation scores indicate closer repertoires.

## Discussion

In this study, we provide a comprehensive characterization of the Treg and Tconv phenotypes and repertoires of WT, non-arthritic SKG and arthritic ZAC mice at the single cell level. We identified two transcriptionally distinct populations of Th17 cells within tissues of arthritic ZAC mice: migratory and pathogenic Th17 cells. Pathogenic Th17 cells have an osteoclastogenic transcriptional signature, are almost exclusively detected in joints and harbor unique and convergent TCRs with a greater similarity to the TCRs of WT Treg cells, which are generally understood to be biased toward self-reactivity(31–34).

It is possible that the pathogenic Th17 population is only acquired within joints by cells harboring self-reactive TCRs, and that these cells are critical for the establishment of disease; whereas migratory Th17 cells may contribute to disease progression and maintenance and represent a prior differentiation state for the cells that later acquire a pathogenic Th17 phenotype. This hypothesis is supported by the observation of a degree of clone sharing between migratory and pathogenic Th17 cells, as well as by trajectory prediction. Our results are also consistent with previous research showing that in experimental autoimmune encephalomyelitis (EAE), Th17 cells progress from a self-renewing cellular, to an effector pre-Th1-like phenotype characterized by the expression of chemokine receptors, to a Th17/Th1-like effector phenotype that exhibits a pathogenic signature with high expression of *Tnfsf11*. Notably, in line with our findings, these cells also display differential enrichment in tissues from lymph nodes to central nervous system (CNS); however, their TCR repertoires were not explored(35).

In a typical immune response, T cells that infiltrate target tissues include both antigen-specific cells and those activated in a bystander manner, recruited by inflammatory signals. In autoimmune diseases, the proportion of tissue-infiltrating T cells that are truly self-reactive remains uncertain. Identifying these self-reactive T cells is essential for understanding the mechanisms driving these disorders and developing more targeted therapeutic strategies. Our study identified a pathogenic subset of joint-infiltrating T cells with a TCR repertoire that is distinctly skewed towards self-reactivity. The selective enrichment of these pathogenic cells in inflamed joints, both in mouse models and human RA synovia, underscores their self-reactive nature and highlights their potential role in recognizing tissue-specific self-antigens.

While other pathogenic Th17 cell subtypes in autoimmune arthritis have been described in the literature(16),(17), the transcriptional profile of the cells identified in our study is distinct, suggesting a unique subset. Although some degree of self-reactivity has been reported for Th17.1 cells(36), the specific TCRs involved, and the characteristics of their repertoire remain largely unexplored. In a separate study, a population of T peripheral helper (Tph) cells, characterized by their expansion and expression of B cell-helper factors, was identified in the joints of seropositive RA patients(37). These cells were later shown to proliferate in autologous mixed lymphocyte reaction (AMLR) assays, indicating self-reactivity(38). However, seropositive RA represents only a subset of RA patients(39), and it is unlikely that the same antigens drive both B cell help and antibody production, as well as local inflammation and tissue destruction. Therefore, our identification of self-reactive cells involved in osteoclastogenesis and tissue destruction is critical for uncovering tissue epitopes relevant across different clinical groups of RA. Furthermore, the single-cell approach in our study enables the retrieval of full and paired-chain TCR sequences, offering the potential for future determination of antigen specificity in the context of a complete TCR repertoire.

In line with this, we observed a number of public clones within ZAC Tconv cells, particularly concentrated within both Th17 phenotypes, and notably expanded. Among the CDR3 sequences of public clones, we identified only one exact match to a TCR in the VDJdb database(40) of TCR sequences with known antigen specificities (Supplementary Table 9). Unexpectedly, the match corresponded to a CD8 TCR, and its target peptide was presented in a class I MHC. Although the presence of “class-mismatched” T cells has been reported in the literature(41–43), it remains unknown whether this phenomenon occurs in this or other self-reactive clones in ZAC mice. Consequently, the identity of any potential epitopes or amino acid motifs remains an open problem.

In addition to these observations on the pathogenic phenotype of ZAC Th17 cells, we carried out the first systematic investigation at single cell resolution of the repertoire shift of conventional T cells upon TCR signal weakening. Our results are in line with the observation that T cells infiltrating prostatic autoimmune lesions in Aire^-/-^ mice express receptors preferentially found in Tregs from Aire^+/+^ mice(44). In addition, exFoxP3 cells display a higher osteoclastogenic potential as compared to Th17 derived from naïve CD4^+^ T cells, presumably due to their higher affinity to self-antigens(17). Together with our findings, these reports point to a key role of TCRs normally occurring in the Treg compartment in the recognition of self-antigens, and suggests that aberrant expression of such TCRs in Tconv cells may drive autoimmune disease.

Finally, the repertoire shift hypothesis raises the prospect of loss of certain TCR specificities within the Treg compartment. Considering that antigen recognition by the Treg TCR appears to be less degenerate than in Tconv cells(29, 45, 46), the loss of certain Treg specificities could profoundly impact tolerance towards specific self-antigens(30). While it is generally acknowledged that Treg and Tconv repertoires are distinct, a small proportion of their repertoires typically overlap(47–49). In this study, we observed a reduction in the similarity between ZAC Treg and Tconv repertoires, specially within activated Tconv cell groups, suggesting a divergence in the Treg-Tconv repertoires that could result in the loss of critical Treg function. Therefore, it is possible to speculate that moderate TCR weakening by ZAP-70 mutation may not only shift the Tconv repertoire but also deplete the Treg repertoire of critical TCRs. The antigen specificity of such TCRs remains unknown, and thus worthy of future study as a starting point for development of TCR-transgenic Treg immunotherapy for the treatment of autoimmune diseases(50).

Taken together, the impact of weakened TCR signaling on the development of autoimmune disease is multifaceted. Firstly, it allows T cells with self-reactive TCRs to escape negative selection or diversion into the Treg lineage, instead differentiating into self-reactive Tconvs. These cells can subsequently acquire a pathogenic phenotype and accumulate in affected tissues. Simultaneously, the Treg repertoire may lose some critical TCRs, leading to both a shift and bifurcation in the repertoire. In addition, there is evidence that Treg effector activity upon TCR weakening is significantly disrupted(5), rendering Tregs insufficient for the suppression of self-reactive Tconvs (Fig. 4).

**Figure 4.**
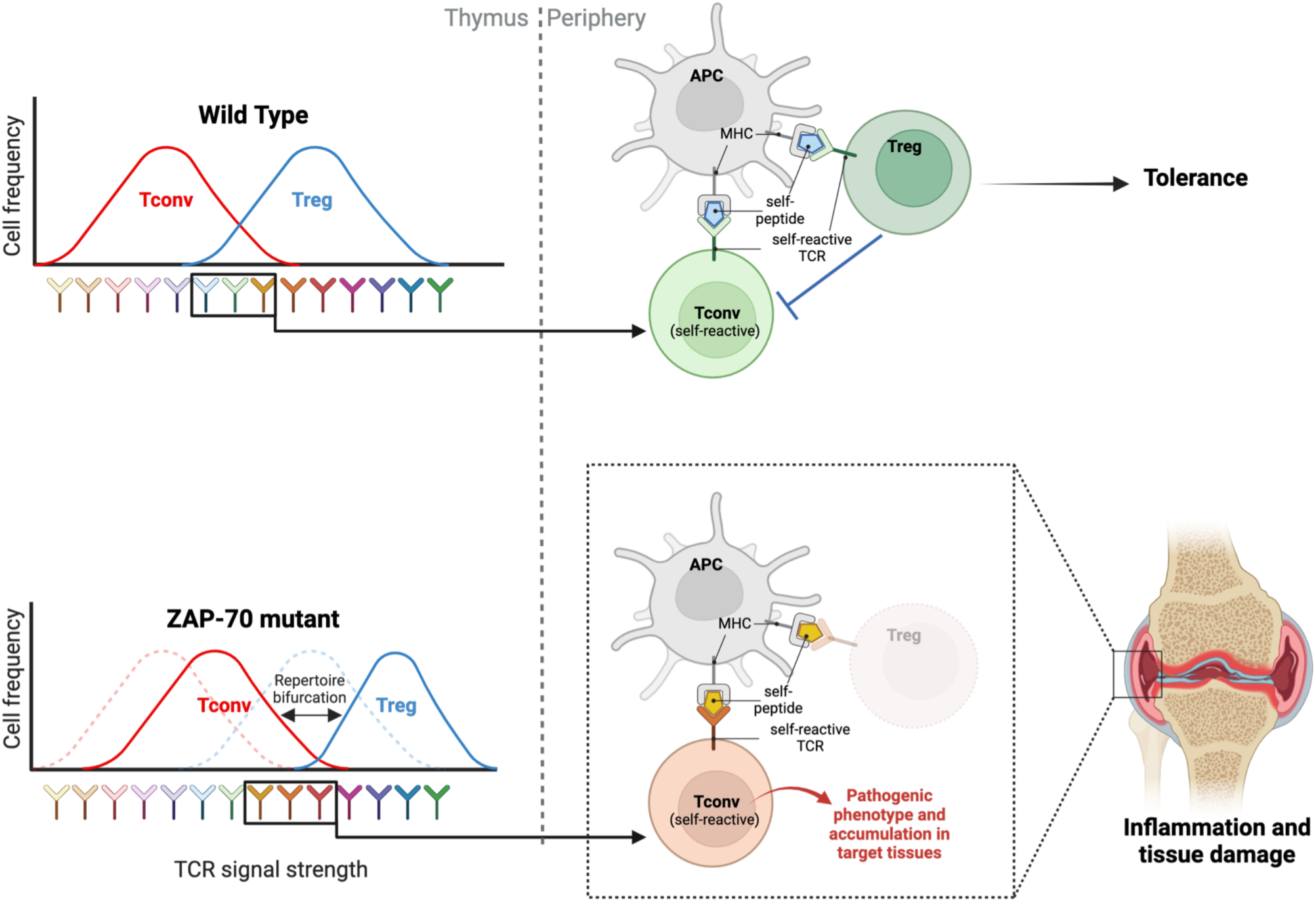
Consequences of weakened TCR signaling in the shaping of Treg and Tconv repertoires and autoimmunity. The SKG (W163C) and ZAC (H165A) mutations in the ZAP-70 gene reduce its affinity for phosphorylated CD3 ζ. As a result, TCR signaling is attenuated. A systemic weakening of TCR signaling would disrupt the T cell selection process by allowing the maturation of self-reactive T cells into the Tconv compartment. Simultaneously, a bifurcation of the Treg-Tconv repertoires is observed, with the consequent loss of certain antigen-specific Treg cells that could effectively suppress autoimmune responses in the periphery. These events together, favor the activation of self-reactive T conv cells, their differentiation into pathogenic phenotypes and their accumulation in target tissues. Created with BioRender.com

It should be noted that our study has several limitations, such as the limited sample size and repertoire coverage that can be practically analyzed by current single cell sequencing approaches. In addition, the pathogenic potential of Th17 cells, and their self-reactivity have not been further experimentally assessed. Moreover, the antigen specificity of public clones could only be predicted, which yielded only a single match to a functionally annotated TCR. Future *in vitro* and *in vivo* experiments will be required to elucidate the specific role and antigen specificities of Th17 cell populations in disease establishment and progression. In spite of these limitations, our research expands the cellular and molecular level understanding of the breakdown in tolerance during autoimmune disease and proposes a straightforward path to discovering TCRs involved in RA.

## Methods

### Sex as a biological variable

The experimental design and analyses in this study were conducted without distinguishing between sexes, and sex was not considered as a biological variable.

### Mice

BALB/c and SKG mice were acquired from CLEA Japan. SKG(8), ZAC(5) and FIG (Foxp3-IRES-GFP knock-in(51)) mice were previously described. In this study, FIG BALB/c WT, FIG SKG and FIG ZAC mice were used and generated as previously reported (5). All animals were maintained in Specific pathogen-free (SPF) facilities.

### Droplet-based single cell RNA sequencing

CD4^+^Foxp3^-^ or Foxp3^+^ cells from FIG BALB/c WT, FIG SKG or FIG ZAC mice were sorted by FACSAria SORP (BD Biosciences). For FIG BALB/c WT and FIG SKG mice, cells were obtained from spleens, processed with the Chromium Single Cell 5’ Library & Gel bead kit (10x Genomics), loaded onto a Chromium Single Cell G Chip, and encapsulated in a Chromium single cell controller (10x Genomics) to generate single-cell gel beads in the emulsion (GEMs) following manufacturer’s protocol. After encapsulation, reverse transcription and cDNA amplification were performed, followed by target enrichment of V(D)J segments and the independent construction of V(D)J and Gene expression libraries as recommended by the manufacturer’s instructions.

For FIG ZAC mice, cells from inflamed joint, draining popliteal lymph node (dLN) and spleen from each replicate were barcoded with TotalSeq hashtag antibodies (BioLegend) and then pooled by tissue and cell type (i.e., Joint Tconv, Joint Treg, etc.) before the generation of GEMs. Next, RNA and V(D)J libraries were produced as described above and an additional library for the barcoded antibodies was prepared. HiSeq3000 and NovaSeq6000 were used for sequencing.

### Single-cell RNAseq data processing

Raw reads were aligned to the mouse reference genome (GRCm38 and GRCm38 VDJ v5.0.0, from 10× Genomics), filtered, and demultiplexed to create gene expression matrices by Cell Ranger (v.5.0). The resulting data was analyzed in R software (v 4.1.2) with Seurat v4.0 package(52, 53).

Where necessary, hashtag demultiplexing was performed by the *HTODemux* function from Seurat and only data from cells classified as singlets was kept for further steps. Next, low-quality data was removed from all datasets, this means cells with unique feature counts less than 200 or over 2500, and cells in which mitochondrial reads represented >5% of reads. Only cells annotated to a single alpha and a single beta chain in the V(D)J result were retained for further analysis. Normalization was performed by the “LogNormalize” method from Seurat and the top 2,000 variable genes were identified in each dataset independently. Batch correction and data integration were performed in combinations of data as follows: 1) Tconvs from ZAC tissues; 2) Tregs from ZAC tissues; 3) Tconvs from spleen (including WT, SKG and ZAC samples); 4) Tregs from spleen (including WT, SKG and ZAC samples). Batch correction was performed with the Canek R package(54) using the intersecting variable genes for the datasets to be integrated in each run.

After integration and QC, the final datasets for analysis consisted of: 6,509 cells for ZAC Tconv, 5,306 cells for ZAC Treg, 24,743 cells for Spleen Tconv and 25,227 cells for Spleen Treg (Supplementary Table 1). Uniform manifold approximation and projection (UMAP) was used to visualize gene expression profiles and clusters from the principal components (PCs) representation of the integrated datasets; the number of PCs were independently selected on each dataset as the elbow suggested in variance plot of PCs. Public single-cell RNAseq data was obtained as count matrices and pre-processed accordingly for QC, normalization, batch correction and data integration (when required).

### Single cell gene expression analysis

#### Clustering and cell-type annotation

The identification of cell clusters based on gene expression data was performed in the PCs representation of integrated datasets with the *FindClusters* function from Seurat using a resolution parameter of 0.5. Clusters were annotated by the examination of expression of canonical Treg or Tconv markers together with the inspection of markers defining each cluster and enriched GO Biological Process Terms (Supplementary Notes 1-4). Additional exclusion of artifactual populations was performed after clustering, including cells lacking expression of CD3 (*Cd3d*, *Cd3e*, and *Cd3g)*, invariant NKT (iNKT) cells, and clusters with high expression of Hemoglobin genes (*Hba-a1, Hba-a2, Hbb-bt, Hbb-bs*).

#### DEG and gene set enrichment analysis

Identification of cluster marker genes was performed with *FindAllMarkers* function in Seurat. Significant markers were considered with a fold change (FC) ≥1.2 and adjusted p value ≤0.05, and were used for gene set enrichment analysis.

When the comparison of two populations was required, DEGs were identified with the *FindMarkers* function with the min.pct parameter set at 0.25. Genes with adjusted p value ≤0.05 where considered significantly changed and were classified as upregulated if FC ≥1.5 or downregulated if FC ≤0.67.

Gene set enrichment analysis was performed with enrichR (v3.1) package(55) using the *GO_Biological_Process_2021* database as reference.

For the generation of ZAC migratory and pathogenic Th17 gene signatures for module score calculation, significant markers were considered if fold change (FC) ≥2 and adjusted p value ≤0.05.

#### Trajectory analysis

Trajectory analyses were performed by using the Slingshot Bioconductor package (v2.10.0)(56) with the PCs representation of integrated datasets and their respective cluster labels as inputs. We set the parameter *start.clus=’naive’* to use the cluster of Naive cells as the starting point of the inferred trajectories.

#### Module score calculation

Calculation of expression levels from Th17 ZAC programs in spleen and public human scRNAseq datasets was performed by the *AddModuleScore* function from Seurat using the specified gene lists or their corresponding human orthologs (Supplementary Table 6) and default parameters.

For public scRNAseq datasets, clustering was performed as described above. Where it was possible to annotate Th17 cells, these were separated in an independent Seurat object and processed for module score calculation and visualization. If Th17 cells were not clearly identified, the whole CD4^+^ T cell clusters were separated in independent Seurat objects for module score calculation.

### Single TCR V(D)J sequencing analysis

#### Clonal expansion and repertoire overlap

Clonal expansion was assessed by the calculation of clonal proportions, then, clones were classified by their size as: small (clonal proportion <1×10^-3^), medium (clonal proportion from 1×10^-3^ – 1×10^-2^) and large (clonal proportion >1×10^-2^). Given that only a small sample size could be recovered for some replicates, which inflates clonal proportions of lowly abundant clones, we labeled singletons (clone size=1) separately.

The Morisita-Horn index is a typical metric used in ecology to calculate the similarity of two ecosystems in terms of species distribution and abundance, and it has been introduced as a metric for repertoire overlap analysis; its values go from 0 to1, where 1 represents a perfect overlap(57). The Morisita-Horn index was calculated between repertoires of each individual replicate, at the paired or single CDR3 amino acid level by the Immunarch (v0.9.0) R package. The median Morisita-Horn indices were compared by a Wilcoxon test using a significance level of 0.05. In addition, clonotype tracking across Treg and Tconv phenotypes was performed by Immunarch.

#### Clonal publicity

Public clones were defined as TCRs sharing paired gene usage and CDR3 amino acid sequences and occurring in more than one subject.

#### TCR distances and repertoire-repertoire correlation

In order to evaluate if the overall features of selected repertoires show convergence, we calculated receptor distances with TCRdist. TCRdist allows for the calculation of a distance measure between TCRs by comparing the CDR1, CDR2 and CDR3 amino acid sequences of single or paired TCR chains(23), then, a neighbor distance metric consisting of the weighted average of the distance of a given TCR to the nearest 25^th^ percentile of the other TCRs in the repertoire can be calculated (weighted neighbor distance). Then, distance distributions of selected repertoires were compared and distributions biased toward lower distances were interpreted as an indication of repertoire convergence. In order to restrict the convergence analysis to the qualitative composition of the repertoires, we used unique clone sequences i.e., clonal counts where not considered. Median distances of Naïve, migratory and pathogenic Th17 cells were compared by a Wilcoxon test with a significance level of 0.05. Finally, the epitope-epitope (or repertoire-repertoire) neighbor distance score correlation metric from the TCRdist algorithm was computed for assessment of intra Treg-Tconv repertoire similarity in pooled WT, SKG and ZAC reperotires. Briefly, a score is assigned to each TCR in one repertoire which is the average distance to the nearest TCRs in a reference repertoire. Then, the sets of scores obtained by two different repertoires can be compared over the entire merged set of TCRs and the Pearson correlation score is calculated. In principle, two similar repertoires will be well correlated.

#### TiRP score calculation

Lagattuta et al proposed a Treg-propensity scoring system for the TCR to quantify the likelihood of Treg cells based on the TCR feature on Treg fate (TiRP score)(24). The source code was obtained from https://github.com/immunogenomics/TiRP. The TiRP score of this study was calculated using the default parameters. For each TCR sequence, the v-gene and amino acid of CDR3 were utilized as input information. Median TiRP scores were compared across groups by a Wilcoxon test with a significance level of 0.05.

#### Published sequence preparation

The published mouse Treg and Tconv TCR beta chain sequence data were obtained from four studies(25–28). The detailed accession ID and references were summarized in Supplementary Table 14. The annotation of raw sequences was performed by mixcr (3.0.13)(58) with align, assemble, assembleContigs, and exportClones process. Sequences with complete frameworks and CDR regions were used in this study.

#### Sequence searching and comparison

We implemented the TCR sequence searching using Interclone method(59). The source code was deposited in https://gitlab.com/sysimm/interclone and the searching parameter using the default setting. In brief, InterClone regards the CDR1,2,3 of one TRB seq as a pseudo sequence and considers one pair of hitting using the following CDR similarity and coverage thresholds: CDR1 ≥ 90%, CDR2 ≥ 90%, CDR3 ≥ 80%, and Coverage ≥ 90%. We searched Tconv and Treg TCR templates from the same reference using identical inputs. Then we compare the corresponding hit numbers with Tconv and Treg templates for each query and calculate the significant Tconv or Treg-biased query sequence using a hypergeometric p-value cutoff of 0.001. We then quantified the total number of matches in terms of the query repertoire size and normalized the results to the matches in the query WT Tconvs (Fig. 4a).

### Statistical analysis

For identification of cluster markers, significant markers were considered with a FC ≥1.2 and adjusted p value ≤0.05. For DEG analysis between two cell populations, genes with adjusted p value ≤0.05 where considered significantly changed and were classified as upregulated if FC ≥1.5 or downregulated if FC ≤0.67. The median Morisita-Horn indices, median distances of Naïve, migratory and pathogenic Th17 repertoires and median TiRP scores were compared by a Wilcoxon test using a significance level of 0.05. For TCR sequence searching and comparison, we calculated the significant Tconv or Treg-biased query sequence using a hypergeometric p-value cutoff of 0.001.

### Study approval

Animal experiments were approved by the institutional review board of Osaka University, and animals were treated in accordance with the institutional guidelines of the Immunology Frontier Research Center, Osaka University.

## Data availability

The datasets generated during the current study have been deposited in the DDBJ Sequence Read Archive database (accession numbers DRA010311, DRA010223), and NCBI Gene Expression Omnibus (accession number GSE180432).

## Author contributions

D.S. and M.L.C. conceived the study. M.L.C., S.T. and A.T. designed experiments. M.L.C., E.L., and A.T performed experiments. M.L.C., M.L.L., D.D., Z.X., performed data analysis. M.L., A.T. and D.S. interpreted data and wrote the paper. S.T. and Z.X. designed custom software for TCRs analysis. D.M. and S.S. supervised the project. All the authors read and approved the final manuscript.

## Acknowledgements

This work was funded by the Japan Agency for Medical Research and Development (AMED), grant number JP223fa627002 and Platform Project for Supporting Drug Discovery and Life Science Research (Basis for Supporting Innovative Drug Discovery and Life Science Research) under JP21am0101108; and by JSPS KAKENHI Grant Numbers JA23H034980 (to D.S.), 20K16286 (to M.L.C.), and 26860331, 17K15723, 22H02920 (to A.T.). M.L.C. received a postdoctoral scholarship from the Mexican National Council of Science and Technology (CONACYT) number 2018-000022-01EXTV-00314 to perform this research.

---

**Supplementary Figure 1. Single-cell transcriptomic analysis of Tconv cells obtained from tissues of arthritic ZAC mice. a)** UMAP plot of integrated Tconv cells obtained from joints, draining lymph node (dLN) and spleen of arthritic ZAC mice. Each panel is colored according to cell source. **b)** Proportion of annotated Tconv cell populations in tissues of each arthritic ZAC mice. **c)** Expression of top 5 gene markers defining each cluster; clusters were subsampled to 100 cells for visualization. **d)** Heatmap of the top 5 enriched GO Terms in each cluster, color scale represents -log10 of adjusted p values. **(e-g)** UMAP plots of Tconv cells from tissues of arthritic ZAC mice colored according to their expression of **e)** inferred pseudotime values **f)** markers of Th17.1 cells: *Ifng*, *Tbx21* and *Cxcr3*; and **g)** markers of exFoxP3 cells: *Sox4*, *Ccr6*, *Ccl20*, *Il23r*, and *Il17re*.

**Supplementary Figure 2. Single-cell transcriptomic and repertoire analysis of Tconv cells obtained from spleens of WT, SKG and ZAC mice.** UMAP plots of integrated Tconv cells obtained from spleens of WT (n=2), SKG (n=2) and arthritic ZAC mice (n=2). Colored by **(a)** cell source; and **(b)** cluster identity. Unsupervised clustering by the Louvain algorithm was applied, and clusters were annotated as described in the Supplementary Notes. **c)** Relative expression of canonical Tconv markers across gene expression clusters. Circle color and intensity represents average gene expression whereas circle size represents the percentage of cells from each cluster expressing the gene. **d)** Proportion of annotated Tconv cell populations in spleens of WT, SKG, and arthritic ZAC mice. **e)** Expression of top 5 gene markers defining each cluster. Clusters were subsampled to 100 cells for visualization. Significant enriched GO Terms in each cluster are depicted to the right, if no GO Terms were significantly enriched, top GO Term is shown and font color is grayed out. **e)** Heatmap of overlap between cell annotations in the ZAC Tconv dataset (columns) and the WT, SKG, ZAC spleen (Spleen Tconv) dataset (rows). Cell colors represent the proportion of ZAC Tconv annotations within Spleen Tconv annotations (i.e., each column sums up to 1). **g)** UMAP plot of integrated Th17 cells annotated in spleens of WT (n=2), SKG (n=2) and arthritic ZAC mice (n=2). Each panel is colored according to cell source. **h)** Distribution of clonal proportion classes across gene expression clusters in spleens of WT, SKG and ZAC Tconv cells.

**Supplementary Figure 3. The pathogenic Th17 gene signature is uniquely detected in CD4^+^ T cells from inflamed synovia of rheumatoid arthritis (RA) patients.** Gene signatures of migratory Th17 and pathogenic Th17 cells were obtained and used for calculation of module scores in public scRNAseq datasets of **a)** Th17 cells obtained from spleens of WT, SKG and ZAC mice (n=2 each), **c)** CD4+ T cells from peripheral blood of RA patients (n=1)(19), **d)** CD4+ T cells from synovia and infrapatellar fat pad (IPFP) tissue of osteoarthritis (OA) patients (n=3)(20), **e)** Circulating Th17 cells from healthy donors (n=3), and patients with autoimmune diseases (AD) other than RA: myasthenia gravis (MG) (n=3), multiple sclerosis (MS) (n=4), systemic lupus erythematosus (SLE) (n=3)(21); **b**) Violin plot of module scores for the gene signatures of migratory Th17 (left) and pathogenic Th17 (right) cells in Th17 cells from spleens of WT, SKG and arthritic ZAC mice.

**Supplementary Figure 4. CD4^+^ T cells from RA synovia expressing the pathogenic Th17 gene signature are different from Th17.1 or exFoxP3 cells.** UMAP plot of CD4^+^ T cells from inflamed synovia of RA patients (n=3)(18), cells are colored according to the density of expression of individual markers of Th17.1 (*Ifng*, *Tbx21* and *Cxcr3*) **(a)** and exFoxP3 cells (*Sox4*, *Ccr6*, *Ccl20*, *Il23r*, and *Il17re*) **(c)**; or as module scores **(b, d)**. The region corresponding to the expression of the pathogenic Th17 gene signature is enclosed by a dashed line in **b** and **d**.

**Supplementary Figure 5. Repertoire analysis of Tconv cells obtained from tissues of arthritic ZAC mice.** Distribution of clonal proportion classes across **(a)** ZAC Tconv gene expression clusters. **(b)** gene expression cluster separated by mouse replicate. **c)** Distribution of neighbor distances to the nearest 25th percentile (methods) in the paired chain (left) and beta chain (right) repertoires of Naïve, migratory Th17 and pathogenic Th17 cells. **d)** Public clones and their clonal proportion classification and distribution across Tconv phenotypes in each mouse replicate

**Supplementary Figure 6. Similarity of TCR repertoires of ZAP-70 mutant Tconv cells, to reference WT Treg and WT Tconv repertoires. a)** Distribution of TCR-intrinsic regulatory potential (TiRP) scores for Tconv and Treg repertoires of WT, SKG and ZAC mice. **b)** Median TiRP score change of Tconv and Treg repertoires of WT, SKG and ZAC mice relative to the WT Tconv repertoire. c) Normalized fraction of query WT Tconv, WT Treg, and SKG and ZAC Tconv repertoires matching with reference WT Tconv (magenta section) or WT Treg repertoires (blue section) obtained from reference datasets(26–28). **d)** Normalized fraction of query WT Tconv, WT Treg, and ZAC migratory and pathogenic Th17 repertoires matching with reference WT Tconv or WT Treg repertoires obtained from reference datasets(26–28). No significant matches to reference WT Treg or WT Tconv in the Ko et al., 2020 dataset were found in pathogenic Th17 cells.

**Supplementary Figure 7. Intra-Treg-Tconv similarity in WT, SKG and ZAC repertoires. a)** Morisita-Horn index calculated at the exact CDR3 amino acid level between the Tconv and Treg compartments of WT, SKG and ZAC mice, split by Tconv phenotype (x axis). **(b-h)** Clonotype tracking plots of intra-Treg-Tconv shared clones across Treg and Tconv gene expression clusters for WT **(b-c)**, SKG **(d-e)**, and ZAC **(f-h)** mice. No Treg-Tconv sharing was detected in ZAC Replicate 3.

## References

1. Pisetsky DS. Pathogenesis of autoimmune disease. Nat Rev Nephrol. 2023;19(8):509–24.

2. Robinson WH, Younis S, Love ZZ, Steinman L, and Lanz TV. Epstein-Barr virus as a potentiator of autoimmune diseases. Nat Rev Rheumatol. 2024.

3. Dou DR, Zhao Y, Belk JA, Zhao Y, Casey KM, Chen DC, et al. Xist ribonucleoproteins promote female sex-biased autoimmunity. Cell. 2024;187(3):733–49 e16.

4. Mori S, Kohyama M, Yasumizu Y, Tada A, Tanzawa K, Shishido T, et al. Neoself-antigens are the primary target for autoreactive T cells in human lupus. Cell. 2024;187(21):6071–87 e20.

5. Tanaka A, Maeda S, Nomura T, Llamas-Covarrubias MA, Tanaka S, Jin L, et al. Construction of a T cell receptor signaling range for spontaneous development of autoimmune disease. J Exp Med. 2023;220(2).

6. Siggs OM, Miosge LA, Yates AL, Kucharska EM, Sheahan D, Brdicka T, et al. Opposing functions of the T cell receptor kinase ZAP-70 in immunity and tolerance differentially titrate in response to nucleotide substitutions. Immunity. 2007;27(6):912–26.

7. Ashby KM, and Hogquist KA. A guide to thymic selection of T cells. Nat Rev Immunol. 2024;24(2):103–17.

8. Sakaguchi N, Takahashi T, Hata H, Nomura T, Tagami T, Yamazaki S, et al. Altered thymic T-cell selection due to a mutation of the ZAP-70 gene causes autoimmune arthritis in mice. Nature. 2003;426(6965):454–60.

9. Tanaka S, Maeda S, Hashimoto M, Fujimori C, Ito Y, Teradaira S, et al. Graded attenuation of TCR signaling elicits distinct autoimmune diseases by altering thymic T cell selection and regulatory T cell function. J Immunol. 2010;185(4):2295–305.

10. Brand DD, Latham KA, and Rosloniec EF. Collagen-induced arthritis. Nat Protoc. 2007;2(5):1269–75.

11. Meyer Zu Horste G, Wu C, Wang C, Cong L, Pawlak M, Lee Y, et al. RBPJ Controls Development of Pathogenic Th17 Cells by Regulating IL-23 Receptor Expression. Cell Rep. 2016;16(2):392–404.

12. Ohl K, Nickel H, Moncrieffe H, Klemm P, Scheufen A, Foll D, et al. The transcription factor CREM drives an inflammatory phenotype of T cells in oligoarticular juvenile idiopathic arthritis. Pediatr Rheumatol Online J. 2018;16(1):39.

13. Glal D, Sudhakar JN, Lu HH, Liu MC, Chiang HY, Liu YC, et al. ATF3 Sustains IL-22-Induced STAT3 Phosphorylation to Maintain Mucosal Immunity Through Inhibiting Phosphatases. Front Immunol. 2018;9:2522.

14. Lin CC, Bradstreet TR, Schwarzkopf EA, Jarjour NN, Chou C, Archambault AS, et al. IL-1-induced Bhlhe40 identifies pathogenic T helper cells in a model of autoimmune neuroinflammation. J Exp Med. 2016;213(2):251–71.

15. DiToro D, Harbour SN, Bando JK, Benavides G, Witte S, Laufer VA, et al. Insulin-Like Growth Factors Are Key Regulators of T Helper 17 Regulatory T Cell Balance in Autoimmunity. Immunity. 2020;52(4):650–67 e10.

16. Basdeo SA, Cluxton D, Sulaimani J, Moran B, Canavan M, Orr C, et al. Ex-Th17 (Nonclassical Th1) Cells Are Functionally Distinct from Classical Th1 and Th17 Cells and Are Not Constrained by Regulatory T Cells. J Immunol. 2017;198(6):2249–59.

17. Komatsu N, Okamoto K, Sawa S, Nakashima T, Oh-hora M, Kodama T, et al. Pathogenic conversion of Foxp3+ T cells into TH17 cells in autoimmune arthritis. Nat Med. 2014;20(1):62–8.

18. Etori K, Tanaka S, Tamura J, Hattori K, Kagami SI, Nakamura J, et al. Fibroblast growth factor receptor 1 as a potential marker of terminal effector peripheral T helper cells in rheumatoid arthritis patients. Rheumatology (Oxford). 2023;62(11):3763–9.

19. Zhang B, Zhang Y, Xiong L, Li Y, Zhang Y, Zhao J, et al. CD127 imprints functional heterogeneity to diversify monocyte responses in inflammatory diseases. J Exp Med. 2022;219(2).

20. Tang S, Yao L, Ruan J, Kang J, Cao Y, Nie X, et al. Single-cell atlas of human infrapatellar fat pad and synovium implicates APOE signaling in osteoarthritis pathology. Sci Transl Med. 2024;16(731):eadf4590.

21. Yasumizu Y, Takeuchi D, Morimoto R, Takeshima Y, Okuno T, Kinoshita M, et al. Single-cell transcriptome landscape of circulating CD4(+) T cell populations in autoimmune diseases. Cell Genom. 2024;4(2):100473.

22. Tong L, Yu H, Huang X, Shen J, Xiao G, Chen L, et al. Current understanding of osteoarthritis pathogenesis and relevant new approaches. Bone Res. 2022;10(1):60.

23. Dash P, Fiore-Gartland AJ, Hertz T, Wang GC, Sharma S, Souquette A, et al. Quantifiable predictive features define epitope-specific T cell receptor repertoires. Nature. 2017;547(7661):89–93.

24. Lagattuta KA, Kang JB, Nathan A, Pauken KE, Jonsson AH, Rao DA, et al. Repertoire analyses reveal T cell antigen receptor sequence features that influence T cell fate. Nat Immunol. 2022;23(3):446–57.

25. Logunova NN, Kriukova VV, Shelyakin PV, Egorov ES, Pereverzeva A, Bozhanova NG, et al. MHC-II alleles shape the CDR3 repertoires of conventional and regulatory naive CD4(+) T cells. Proc Natl Acad Sci U S A. 2020;117(24):13659–69.

26. Lu DR, Wu H, Driver I, Ingersoll S, Sohn S, Wang S, et al. Dynamic changes in the regulatory T-cell heterogeneity and function by murine IL-2 mutein. Life Sci Alliance. 2020;3(5).

27. Ko A, Watanabe M, Nguyen T, Shi A, Achour A, Zhang B, et al. TCR Repertoires of Thymic Conventional and Regulatory T Cells: Identification and Characterization of Both Unique and Shared TCR Sequences. J Immunol. 2020;204(4):858–67.

28. Wolf KJ, Emerson RO, Pingel J, Buller RM, and DiPaolo RJ. Conventional and Regulatory CD4+ T Cells That Share Identical TCRs Are Derived from Common Clones. PLoS One. 2016;11(4):e0153705.

29. Pohar J, Simon Q, and Fillatreau S. Antigen-Specificity in the Thymic Development and Peripheral Activity of CD4(+)FOXP3(+) T Regulatory Cells. Front Immunol. 2018;9:1701.

30. Ooi JD, Petersen J, Tan YH, Huynh M, Willett ZJ, Ramarathinam SH, et al. Dominant protection from HLA-linked autoimmunity by antigen-specific regulatory T cells. Nature. 2017;545(7653):243–7.

31. Jordan MS, Boesteanu A, Reed AJ, Petrone AL, Holenbeck AE, Lerman MA, et al. Thymic selection of CD4+CD25+ regulatory T cells induced by an agonist self-peptide. Nat Immunol. 2001;2(4):301–6.

32. Hsieh CS, Zheng Y, Liang Y, Fontenot JD, and Rudensky AY. An intersection between the self-reactive regulatory and nonregulatory T cell receptor repertoires. Nat Immunol. 2006;7(4):401–10.

33. Aschenbrenner K, D’Cruz LM, Vollmann EH, Hinterberger M, Emmerich J, Swee LK, et al. Selection of Foxp3+ regulatory T cells specific for self antigen expressed and presented by Aire+ medullary thymic epithelial cells. Nat Immunol. 2007;8(4):351–8.

34. Lee HM, Bautista JL, Scott-Browne J, Mohan JF, and Hsieh CS. A broad range of self-reactivity drives thymic regulatory T cell selection to limit responses to self. Immunity. 2012;37(3):475–86.

35. Gaublomme JT, Yosef N, Lee Y, Gertner RS, Yang LV, Wu C, et al. Single-Cell Genomics Unveils Critical Regulators of Th17 Cell Pathogenicity. Cell. 2015;163(6):1400–12.

36. Paroni M, Maltese V, De Simone M, Ranzani V, Larghi P, Fenoglio C, et al. Recognition of viral and self-antigens by T(H)1 and T(H)1/T(H)17 central memory cells in patients with multiple sclerosis reveals distinct roles in immune surveillance and relapses. J Allergy Clin Immunol. 2017;140(3):797–808.

37. Rao DA, Gurish MF, Marshall JL, Slowikowski K, Fonseka CY, Liu Y, et al. Pathologically expanded peripheral T helper cell subset drives B cells in rheumatoid arthritis. Nature. 2017;542(7639):110–4.

38. Sakuragi T, Yamada H, Haraguchi A, Kai K, Fukushi JI, Ikemura S, et al. Autoreactivity of Peripheral Helper T Cells in the Joints of Rheumatoid Arthritis. J Immunol. 2021;206(9):2045–51.

39. Cunningham KY, Hur B, Gupta VK, Arment CA, Wright KA, Mason TG, et al. Patients with ACPA-positive and ACPA-negative rheumatoid arthritis show different serological autoantibody repertoires and autoantibody associations with disease activity. Sci Rep. 2023;13(1):5360.

40. Shugay M, Bagaev DV, Zvyagin IV, Vroomans RM, Crawford JC, Dolton G, et al. VDJdb: a curated database of T-cell receptor sequences with known antigen specificity. Nucleic Acids Res. 2018;46(D1):D419–D27.

41. Boyle LH, Goodall JC, and Gaston JS. Major histocompatibility complex class I-restricted alloreactive CD4+ T cells. Immunology. 2004;112(1):54–63.

42. Boyle LH, Goodall JC, Opat SS, and Gaston JS. The recognition of HLA-B27 by human CD4(+) T lymphocytes. J Immunol. 2001;167(5):2619–24.

43. Singh NK, Alonso JA, Devlin JR, Keller GLJ, Gray GI, Chiranjivi AK, et al. A class-mismatched TCR bypasses MHC restriction via an unorthodox but fully functional binding geometry. Nat Commun. 2022;13(1):7189.

44. Malchow S, Leventhal DS, Lee V, Nishi S, Socci ND, and Savage PA. Aire Enforces Immune Tolerance by Directing Autoreactive T Cells into the Regulatory T Cell Lineage. Immunity. 2016;44(5):1102–13.

45. Kieback E, Hilgenberg E, Stervbo U, Lampropoulou V, Shen P, Bunse M, et al. Thymus-Derived Regulatory T Cells Are Positively Selected on Natural Self-Antigen through Cognate Interactions of High Functional Avidity. Immunity. 2016;44(5):1114–26.

46. Leonard JD, Gilmore DC, Dileepan T, Nawrocka WI, Chao JL, Schoenbach MH, et al. Identification of Natural Regulatory T Cell Epitopes Reveals Convergence on a Dominant Autoantigen. Immunity. 2017;47(1):107–17 e8.

47. Hsieh CS, Liang Y, Tyznik AJ, Self SG, Liggitt D, and Rudensky AY. Recognition of the peripheral self by naturally arising CD25+ CD4+ T cell receptors. Immunity. 2004;21(2):267–77.

48. Pacholczyk R, Ignatowicz H, Kraj P, and Ignatowicz L. Origin and T cell receptor diversity of Foxp3+CD4+CD25+ T cells. Immunity. 2006;25(2):249–59.

49. Al Khabouri S, Benson RA, Prendergast CT, Gray JI, Otto TD, Brewer JM, et al. TCRbeta Sequencing Reveals Spatial and Temporal Evolution of Clonal CD4 T Cell Responses in a Breach of Tolerance Model of Inflammatory Arthritis. Front Immunol. 2021;12:669856.

50. Eggenhuizen PJ, Cheong RMY, Lo C, Chang J, Ng BH, Ting YT, et al. Smith-specific regulatory T cells halt the progression of lupus nephritis. Nat Commun. 2024;15(1):899.

51. Ohkura N, Hamaguchi M, Morikawa H, Sugimura K, Tanaka A, Ito Y, et al. T cell receptor stimulation-induced epigenetic changes and Foxp3 expression are independent and complementary events required for Treg cell development. Immunity. 2012;37(5):785–99.

52. Satija R, Farrell JA, Gennert D, Schier AF, and Regev A. Spatial reconstruction of single-cell gene expression data. Nat Biotechnol. 2015;33(5):495–502.

53. Stoeckius M, Zheng S, Houck-Loomis B, Hao S, Yeung BZ, Mauck WM, 3rd, et al. Cell Hashing with barcoded antibodies enables multiplexing and doublet detection for single cell genomics. Genome Biol. 2018;19(1):224.

54. Loza M, Teraguchi S, Standley DM, and Diez D. Unbiased integration of single cell transcriptome replicates. NAR Genom Bioinform. 2022;4(1):lqac022.

55. Chen EY, Tan CM, Kou Y, Duan Q, Wang Z, Meirelles GV, et al. Enrichr: interactive and collaborative HTML5 gene list enrichment analysis tool. BMC Bioinformatics. 2013;14:128.

56. Street K, Risso D, Fletcher RB, Das D, Ngai J, Yosef N, et al. Slingshot: cell lineage and pseudotime inference for single-cell transcriptomics. BMC Genomics. 2018;19(1):477.

57. Chiffelle J, Genolet R, Perez MA, Coukos G, Zoete V, and Harari A. T-cell repertoire analysis and metrics of diversity and clonality. Curr Opin Biotechnol. 2020;65:284–95.

58. Bolotin DA, Poslavsky S, Mitrophanov I, Shugay M, Mamedov IZ, Putintseva EV, et al. MiXCR: software for comprehensive adaptive immunity profiling. Nat Methods. 2015;12(5):380–1.

59. Wilamowski J, Xu Z, Ismanto HS, Li S, Teraguchi S, Llamas-Covarrubias MA, et al. InterClone: Store, Search and Cluster Adaptive Immune Receptor Repertoires. bioRxiv. 2022:2022.07.31.501809.

