## Supplementary Figures for "The TCRs of pathogenic Th17 cells in arthritogenic mice are shifted toward a Treg-like repertoire"

### Supplementary figure 1

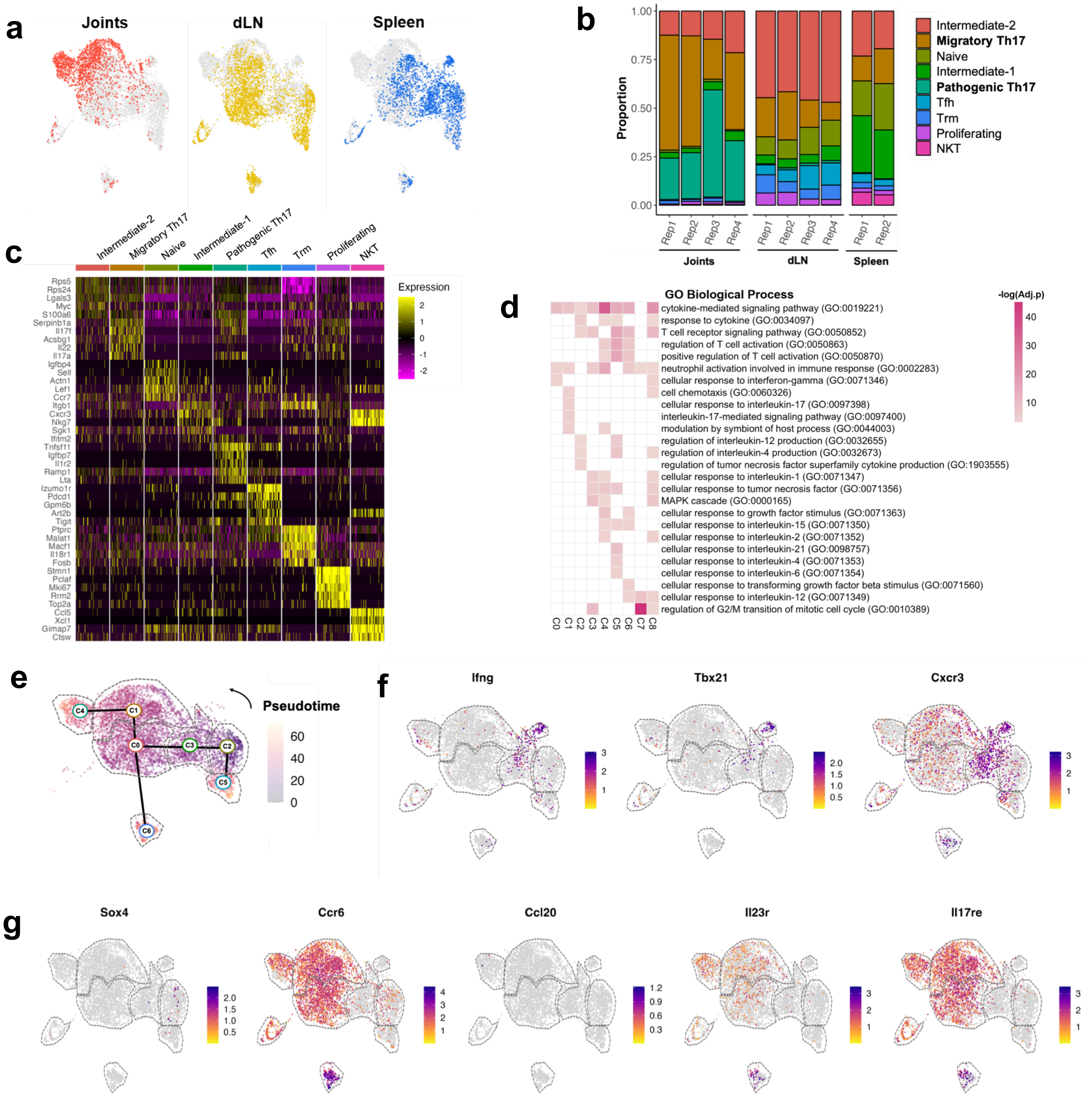


**Supplementary Figure 1. Single-cell transcriptomic analysis of Tconv cells obtained from tissues of arthritic ZAC mice. a)** UMAP plot of integrated Tconv cells obtained from joints, draining lymph node (dLN) and spleen of arthritic ZAC mice. Each panel is colored according to cell source. **b)** Proportion of annotated Tconv cell populations in tissues of each arthritic ZAC mice. **c)** Expression of top 5 gene markers defining each cluster; clusters were subsampled to 100 cells for visualization. **d)** Heatmap of the top 5 enriched GO Terms in each cluster, color scale represents -log10 of adjusted p values. UMAP plots of Tconv cells from tissues of arthritic ZAC mice colored according to their expression of **(e)** inferred pseudotime values **(f)** markers of Th17.1 cells: *Ifng*, *Tbx21* and *Cxcr3*; and **(g)** markers of exFoxP3 cells: *Sox4*, *Ccr6*, *Ccl20*, *Il23r*, and *Il17re.*

### Supplementary figure 2

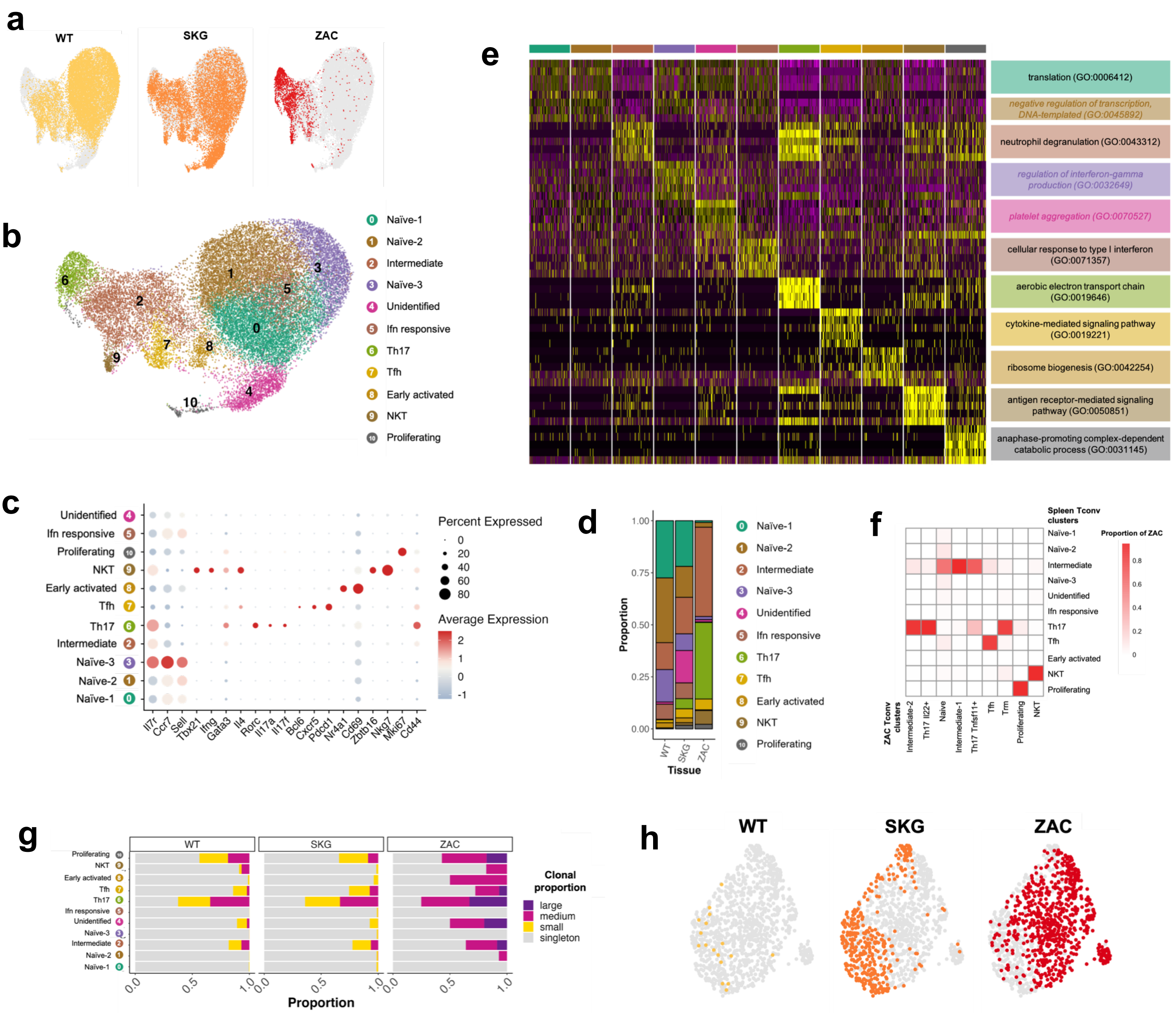


**Supplementary Figure 2. Single-cell transcriptomic and repertoire analysis of Tconv cells obtained from spleens of WT, SKG and ZAC mice.** UMAP plots of integrated Tconv cells obtained from spleens of WT (n=2), SKG (n=2) and arthritic ZAC mice (n=2) colored by **(a)** cell source; and **(b)** cluster identity. Unsupervised clustering by the Louvain algorithm was applied, and clusters were annotated as described in the Supplementary Notes. **c)** Relative expression of canonical Tconv markers across gene expression clusters. Circle color and intensity represents average gene expression whereas circle size represents the percentage of cells from each cluster expressing the gene. **d)** Proportion of annotated Tconv cell populations in spleens of WT, SKG, and arthritic ZAC mice. **e)** Expression of top 5 gene markers defining each cluster. Clusters were subsampled to 100 cells for visualization. Significant enriched GO Terms in each cluster are depicted to the right, if no GO Terms were significantly enriched, top GO Term is shown and font color is grayed out. **f)** Heatmap of overlap between cell annotations in the ZAC Tconv dataset (columns) and the WT, SKG, ZAC spleen (Spleen Tconv) dataset (rows). Cell colors represent the proportion of ZAC Tconv annotations within Spleen Tconv annotations (i.e., each column sums up to 1). **g)** Distribution of clonal proportion classes across gene expression clusters in spleens of WT, SKG and ZAC Tconv cells. **h)** UMAP plot of integrated Th17 cells annotated in spleens of WT (n=2), SKG (n=2) and arthritic ZAC mice (n=2). Each panel is colored according to cell source.

### Supplementary figure 3

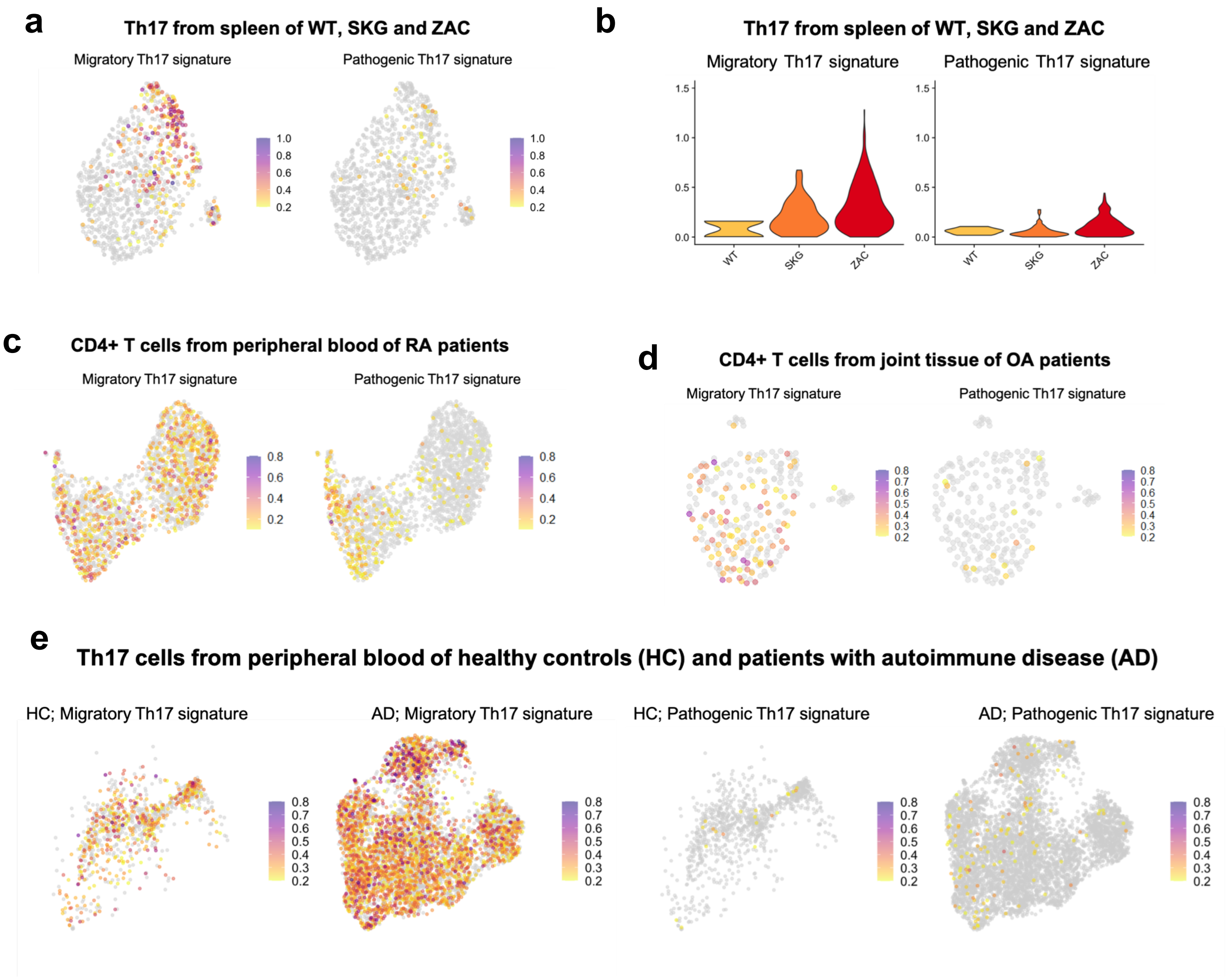


**Supplementary Figure 3. The pathogenic Th17 gene signature is uniquely detected in CD4+ T cells from inflamed synovia of rheumatoid arthritis (RA) patients.** Gene signatures of migratory Th17 and pathogenic Th17 cells were obtained and used for calculation of module scores in public scRNAseq datasets of **a)** Th17 cells obtained from spleens of WT, SKG and ZAC mice (n=2 each), **c)** CD4+ T cells from peripheral blood of RA patients (n=1)[19], **d)** CD4+ T cells from synovia and infrapatellar fat pad (IPFP) tissue of osteoarthritis (OA) patients (n=3)[20], **e)** Circulating Th17 cells from healthy donors (n=3), and patients with autoimmune diseases (AD) other than RA: myasthenia gravis (MG) (n=3), multiple sclerosis (MS) (n=4), systemic lupus erythematosus (SLE) (n=3)[21]; **b)** Violin plot of module scores for the gene signatures of migratory Th17 (left) and pathogenic Th17 (right) cells in Th17 cells from spleens of WT, SKG and arthritic ZAC mice.

### Supplementary figure 4

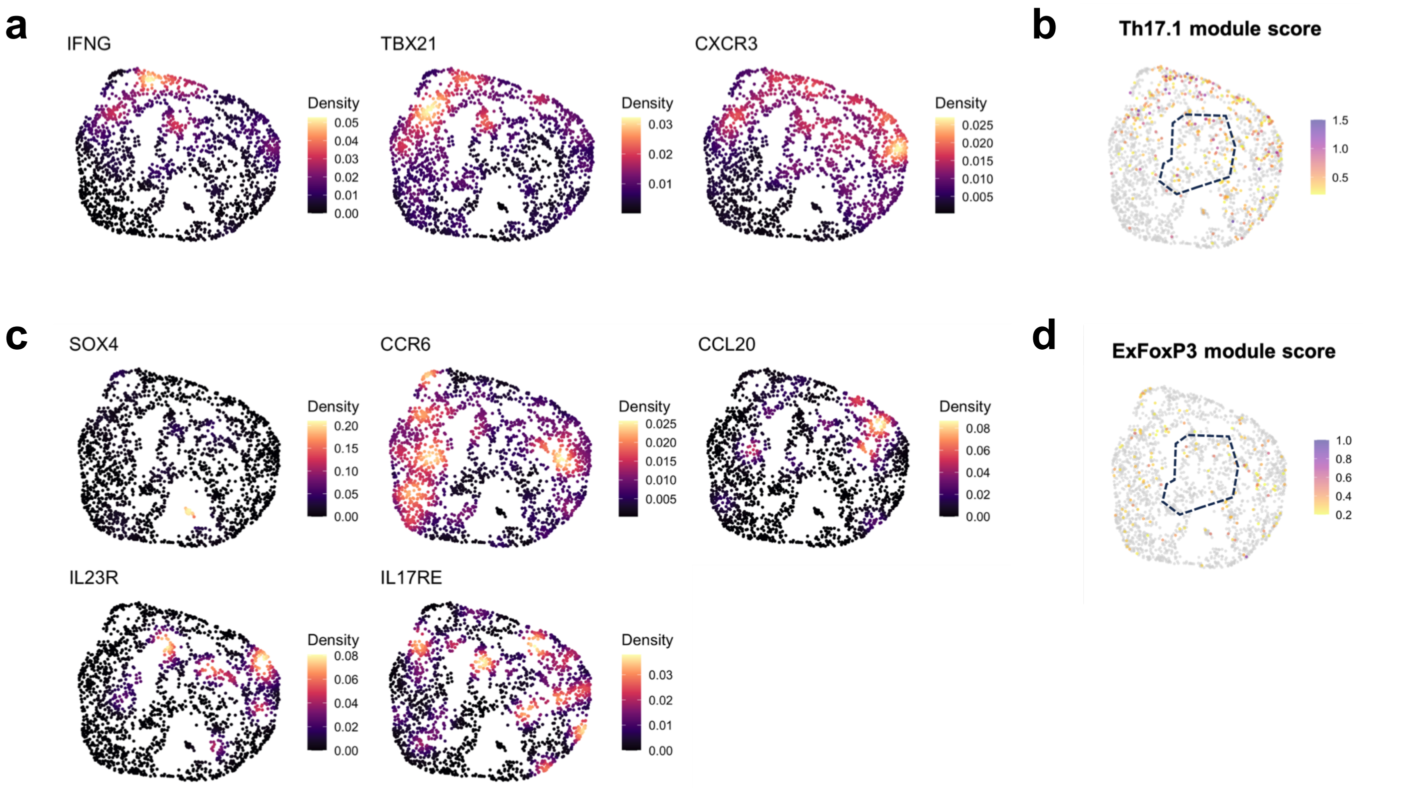


**Supplementary Figure 4. CD4+ T cells from RA synovia expressing the pathogenic Th17 gene signature are different from Th17.1 or exFoxP3 cells.** UMAP plot of CD4+ T cells from inflamed synovia of RA patients (n=3)[18], cells are colored according to the density of expression of individual markers of Th17.1 (*Ifng, Tbx21* and *Cxcr3*) **(a)** and exFoxP3 cells (*Sox4*, *Ccr6*, *Ccl20*, *Il23r*, and *Il17re*) **(c)**; or as module scores **(b, d)**. The region corresponding to the expression of the pathogenic Th17 gene signature is enclosed by a dashed line in **b** and **d**.

### Supplementary figure 5

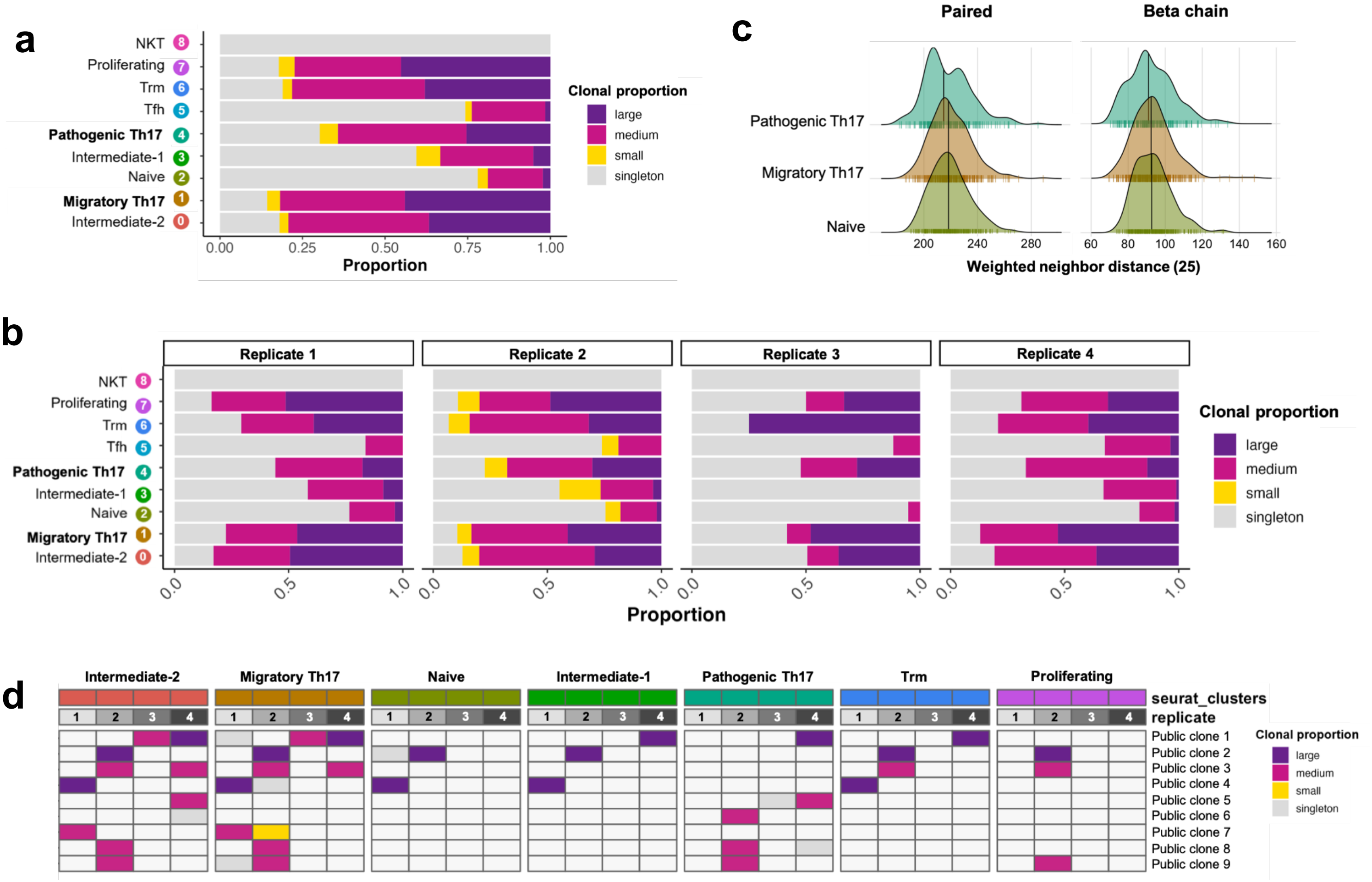


**Supplementary Figure 5. Repertoire analysis of Tconv cells obtained from tissues of arthritic ZAC mice.** Distribution of clonal proportion classes across **(a)** ZAC Tconv gene expression clusters. **(b)** gene expression cluster separated by mouse replicate. **c)** Distribution of neighbor distances to the nearest 25th percentile (methods) in the paired chain (left) and beta chain (right) repertoires of Naïve, migratory Th17 and pathogenic Th17 cells. **d)** Public clones and their clonal proportion classification and distribution across Tconv phenotypes in each mouse replicate

### Supplementary Figure 6

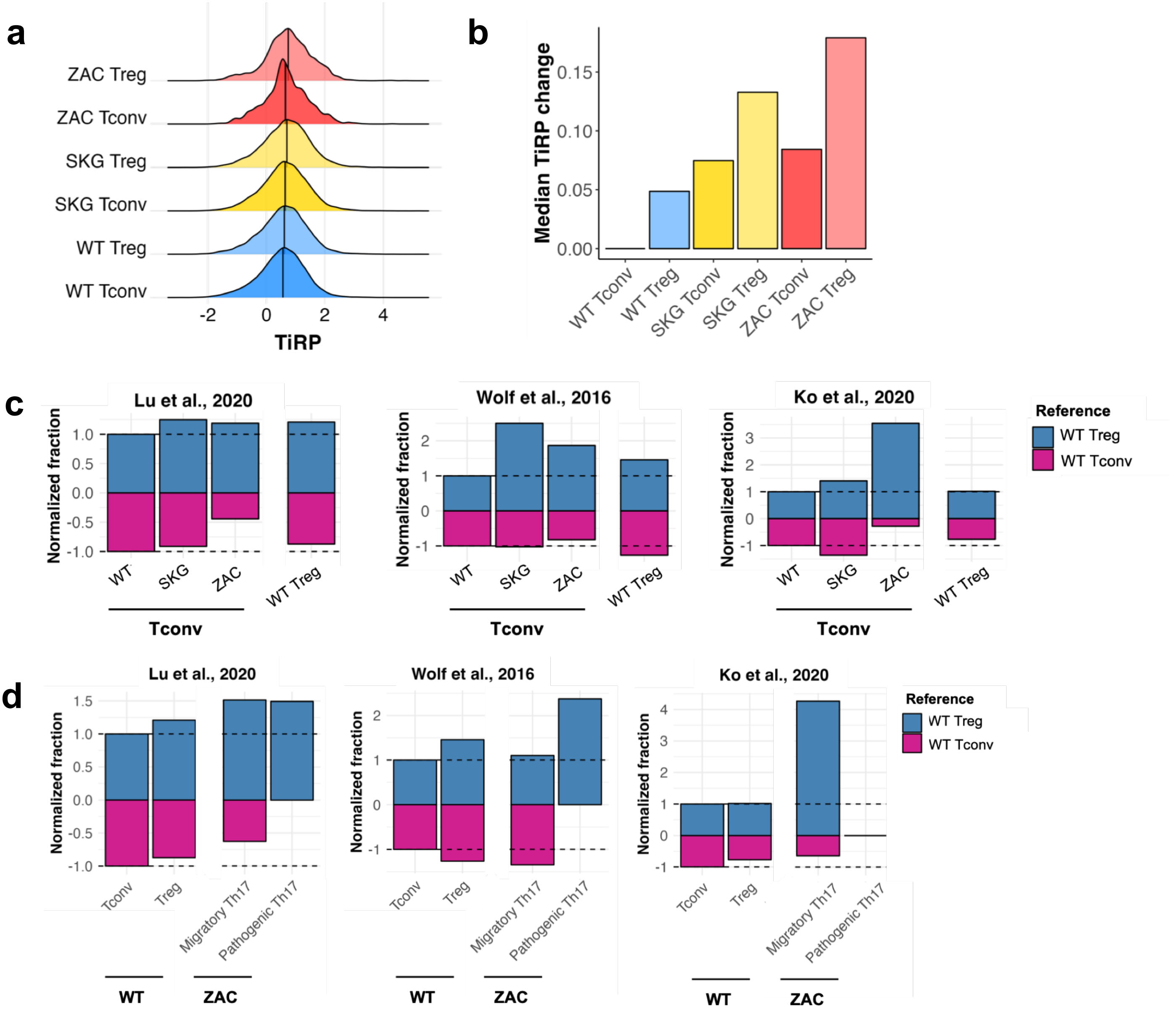


**Supplementary Figure 6. Similarity of TCR repertoires of ZAP-70 mutant Tconv cells, to reference WT Treg and WT Tconv repertoires. a)** Distribution of TCR-intrinsic regulatory potential (TiRP) scores for Tconv and Treg repertoires of WT, SKG and ZAC mice. **b)** Median TiRP score change of Tconv and Treg repertoires of WT, SKG and ZAC mice relative to the WT Tconv repertoire. **c)** Normalized fraction of query WT Tconv, WT Treg, and SKG and ZAC Tconv repertoires matching with reference WT Tconv (magenta section) or WT Treg repertoires (blue section) obtained from reference datasets[26-28]. **d)** Normalized fraction of query WT Tconv, WT Treg, and ZAC migratory and pathogenic Th17 repertoires matching with reference WT Tconv or WT Treg repertoires obtained from reference datasets[26-28]. No significant matches to reference WT Treg or WT Tconv in the Ko et al., 2020 dataset were found in pathogenic Th17 cells.

### Supplementary Figure 7

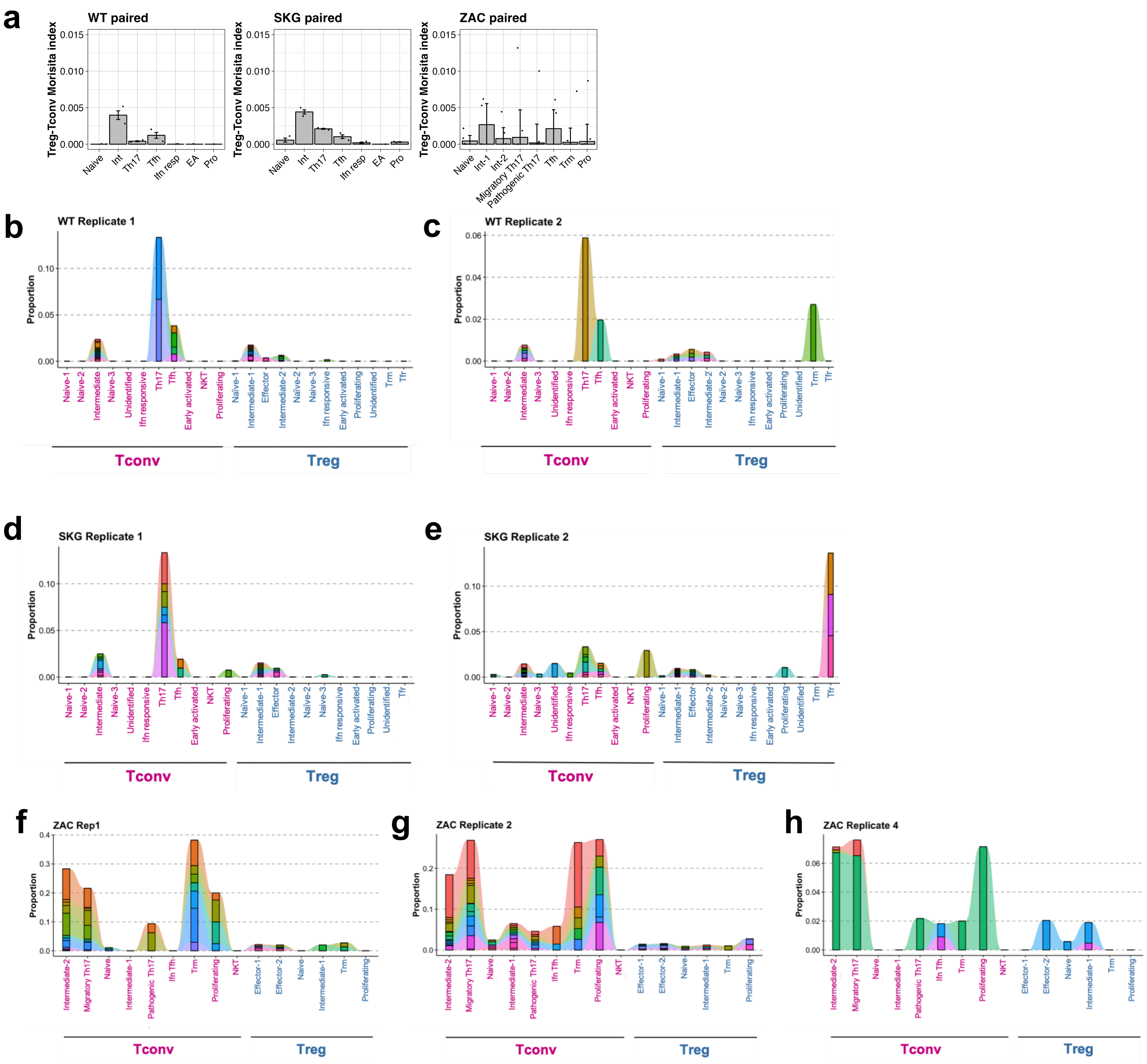


**Supplementary Figure 7. Intra-Treg-Tconv similarity in WT, SKG and ZAC repertoires. a)** Morisita-Horn index calculated at the exact CDR3 amino acid level between the Tconv and Treg compartments of WT, SKG and ZAC mice, split by Tconv phenotype (x axis). **(b-h)** Clonotype tracking plots of intra-Treg-Tconv shared clones across Treg and Tconv gene expression clusters for WT **(b-c)**, SKG **(d-e)**, and ZAC **(f-h)** mice. No Treg-Tconv sharing was detected in ZAC Replicate 3.
