## Supplementary Notes for "The TCRs of pathogenic Th17 cells in arthritogenic mice are shifted toward a Treg-like repertoire"

**Supplementary Note 1**

T cell canonical gene markers, differentially expressed genes (DEG) defining each cluster, and enrichment of Gene Ontology (GO) terms (Fig. 1e, Supplementary Fig. 1c-d, Supplementary Table 2-3) were used for cluster annotation. Expression of *Sell*, *Ccr7*, and *Il7r* indicated the existence of Naïve cells (cluster 2), which accounted for about 20% of spleen cells and 11% of draining lymph node (dLN) cells but less than 1% of joint Tconvs. The expression of *Bcl6, Ccr5, Pdcd1,* together with *Izumo1r* indicated the presence of follicular helper T cells (Tfh, cluster 5) that were abundant in dLN. High expression of *Il7r, Cd44, Cd69* and *Itgae* (encoding CD103), consistent with a tissue-resident memory T (Trm) cells was observed in cluster 6 which occurred in all tissues. Expression of *Mki67* and *Top2a*, indicated proliferative lymphocytes (cluster 7). Gene markers *Zbtb16, Nkg7,* and *Ccl5* indicated typical NKT cells (cluster 8) that were consistently enriched in the spleen. We could not detect a clear pattern of canonical T cell markers of activation in clusters 0 and 3 (Fig. 1e). Cluster 3 was more abundant in spleen (~27%) (Fig. 1d), and showed some degree of overlap with the transcriptional profile of Naïve cells (Supplementary Fig. 1c), but cluster 3 differentially expresses *Cxcr3*, suggesting that these cells are undergoing an activation process. In addition, the inference of cell pseudotimes and trajectory prediction, positioned cluster 3 cells in an intermediate step in trajectory, just after the naïve state (Supplementary Fig. 1e). Thus, we annotated cluster 3 as Intermediate-1 cells. In contrast, cluster 0, which is abundant in dLN (~45%), showed partial overlap with the transcriptional signature of Th17 clusters (Supplementary Fig. 1c), and trajectory inference positioned cluster 0 as an intermediate step just prior to the Th17 states (Supplementary Fig. 1e), thus, we labeled cluster 0 as Intermediate-2.

**Supplementary Note 2**

UMAP visualization revealed a considerable overlap between WT and SKG Tconv transcriptional profiles compared to ZAC mice (Supplementary Fig. 2a). Unsupervised clustering delineated a total of eleven Spleen Tconv clusters which included three naïve groups (Naive-1, Naive-2 and Naive-3: *Sell, Il7r, Ccr7*), Th17 (*Rorc, Il17a, Il17f*), Tfh (*Bcl6, Cxcr5, Pdcd1*), early activated (*Nr4a1*, *Cd69)*, NKT (*Nkg7*, *Zbtb16*), and proliferating (*Mki67*) (Supplementary Fig. 2b-c). Two clusters (2, 5) could not be clearly assigned to a T cell subtype by simple examination of canonical markers and were annotated as Intermediate, and Interferon responsive (*Stat1*, *Gbp2*, *Gbp4*), based on trajectory analysis (not shown) and enrichment analysis of GO Terms (Supplementary Fig. 2e). Finally, one group of cells (cluster

4) showed low expression of all the analyzed canonical markers, no significantly enriched GO Terms, and was represented in only one replicate (Supplementary Fig. 2c, 2e) so it was attributed to a batch effect and denoted as Unidentified. Importantly, annotations for the spleen datasets were mostly consistent with the annotations assigned in the ZAC dataset alone (Supplementary Fig. 2f). As expected, Th17 cells were clonally expanded in all mice, particularly in ZAC (Supplementary Fig. 2g).
